# Single-molecule tracking of Nodal and Lefty in live zebrafish embryos supports hindered diffusion model

**DOI:** 10.1101/2022.04.05.487143

**Authors:** Timo Kuhn, Amit N. Landge, David Mörsdorf, Jonas Coßmann, Johanna Gerstenecker, Patrick Müller, J. Christof M. Gebhardt

## Abstract

The influential hindered diffusion model postulates that the global movement of a signaling molecule through an embryo is affected by local tissue geometry and binding-mediated hindrance, but these effects have not been directly demonstrated *in vivo* for any signaling molecule. Nodal and Lefty are a prime example of an activator-inhibitor signaling pair whose different global diffusivities are thought to arise from differential hindrance. Here, we used single-molecule tracking of Nodal and Lefty to directly probe the tenets of the hindered diffusion model on the nanoscale. We visualized individual fluorescently-tagged Nodal and Lefty molecules in developing zebrafish embryos using reflected light-sheet microscopy. Single-particle tracking revealed molecules in three states: molecules diffusing in extracellular cavities, molecules diffusing within cell-cell interfaces, and molecules bound to cell membranes. While the diffusion coefficients of molecules were high in extracellular cavities, mobility was reduced and bound fractions were higher within cell-cell interfaces; counterintuitively, molecules nevertheless accumulated in cavities. Using agent-based simulations, we identified the geometry of the extracellular space as a key factor influencing the accumulation of molecules in cavities. For Nodal, the fraction of molecules in the bound state was larger than for Lefty, and individual Nodal molecules had binding times of tens of seconds. Together, our single-molecule measurements and simulations provide direct support for the hindered diffusion model in a developing embryo and yield unprecedented insights into the nanometer to micrometer scale transport mechanisms that together lead to macroscopic signal dispersal and gradient formation.

## Introduction

The development of an embryo from a single cell to a complex organism is coordinated by cellular communication via signaling molecules called morphogens. Morphogens are produced in localized sources, from which they spread to form concentration gradients. Target cells along a morphogen gradient perceive different amounts and durations of morphogen signaling and respond by switching on different cell fate programs. By coupling molecular concentrations to distributions in space, morphogens can therefore provide positional information to orchestrate tissue patterning^1^.

The range of a morphogen gradient needs to span multiple cell diameters from the source in order to provide positional information. While special transport mechanisms – for instance along cell extensions – are important in certain developmental contexts^2–6^, the most prominent theory to explain the establishment of a morphogen gradient is the synthesis-diffusion-clearance model^7–14^. In this model, morphogens are produced in a localized source, from which they spread into neighboring tissues by diffusion. The length-scale of the gradient is determined by morphogen clearance – degradation or cellular uptake – as well as the morphogen’s diffusivity. While the free diffusivity of a morphogen is a biophysical property that can be influenced by factors in the tissue environment such as temperature and viscosity, the hindered diffusion model postulates that the effective diffusivity of a molecule can be further influenced by transient binding interactions^2,7^. Indeed, there are numerous reports demonstrating direct binding of morphogens to intra- and extracellular molecules such as receptors^14–16^, collagen^17^ and heparin sulphate proteoglycans^7,18–23^ that can modulate the shape of a morphogen gradient, but it remains unclear whether binding truly affects morphogen diffusivity or rather retention, uptake and stability. Beyond flat tissue culture systems^24^, the tenets of the hindered diffusion model – i) free diffusion far away from cell surfaces, ii) hindered diffusion due to the tissue architecture, iii) further reduction due to binding^2,7^– have not been directly demonstrated for any morphogen in an *in vivo* tissue context.

The secreted TGF-β superfamily ligands Nodal and Lefty are prime examples of an activator-inhibitor morphogen pair whose different signaling ranges have been postulated to arise from differential hindrance^2,25,26^. This system has been best characterized in zebrafish embryos, where the Nodal signaling proteins Squint and Cyclops are produced in the marginal zone and induce the formation of mesoderm and endoderm during early development, beginning around 4 h post-fertilization (hpf)^9,27–31^. Nodal signaling is antagonized by secreted Leftys^28,32– 35^, which inhibit Nodal from binding to their receptors^36,37^.

Hindered diffusion has been proposed to result in the formation of Nodal and Lefty concentration gradients^2,26^ where Cyclops has an ultra-short range of only a few micrometers, Squint has a short-to-mid range, Lefty1 acts at a long range, and Lefty2 has an ultra-long range leading to a nearly uniform distribution throughout the embryo^26^. Previous observations of the Nodal/Lefty system are consistent with the hindered diffusion model. First, free diffusion coefficients measured by fluorescence correlation spectroscopy (FCS) in a diffraction-limited spot far away from cell surface yielded similar local diffusion coefficients on a sub-micrometer scale for zebrafish Nodals and Leftys^15,26^. Second, effective diffusion coefficients on a tissue level across a cube of approximately 8 × 8 × 8 cells measured by fluorescence recovery after photobleaching (FRAP) were found to be much lower for Nodals than for Leftys (Cyclops < Squint < Lefty1 < Lefty2)^26,38–40^. Third, it has been shown that Nodals bind their receptors with nanomolar affinity^15^, and manipulating the levels of the co-receptor Oep modulated the Nodal signaling range^41,42^. However, it remains unclear whether and how such treatments affect Nodal and Lefty movement, how effective diffusivity through a tissue emerges from interactions at the molecular scale, how binding on the cell surface contributes to Nodal and Lefty movement, and how tissue geometry affects morphogen spreading.

Here, we present single-molecule imaging and tracking of HaloTag-tagged fluorescent Cyclops, Squint, Lefty1 and Lefty2 in the extracellular environment of live developing zebrafish embryos. We monitored the movement of these morphogens on the nanoscale and observed a major influence of the local extracellular architecture on the diffusion properties. We found that molecules moving in extracellular cavities between cells were predominantly diffusing freely. In contrast, we observed hindered diffusion within cell-cell interfaces with larger bound fractions of Nodal molecules compared to Lefty. Time-lapse microscopy enabled us to observe individual binding events of tens of seconds for Cyclops and Squint. We developed an agent-based model of single-molecule movements and found a major contribution of tissue architecture, receptor levels and affinity on morphogen distributions. Overall, our single-molecule fluorescence measurements directly support a model of hindered diffusion for Nodal and Lefty, where Nodals – but not Leftys – are transiently trapped on the cell surface, explaining their short action range.

## Results

### Single-molecule imaging of HaloTag-labeled morphogens in live zebrafish embryos

To observe the movement of individual morphogens, we used a reflected light-sheet microscope (RLSM), which is ideally suited to image single molecules in live zebrafish embryos^43^. In order to visualize Nodals and Leftys we fused them to HaloTags. We inserted the HaloTag between the pro- and mature domains of Cyclops and Squint and added them to the C-termini of Lefty1 and Lefty2, generating active and properly localized proteins analogous to previous approaches^26^ (Figure 1a, Supplementary Figure 1, Materials and Methods). The HaloTag allows precise titration of the amount of fluorescence label, ensuring low densities of labeled molecules in every frame over the entire measurement period (Supplementary Figure 2). To visualize single molecules, we injected embryos at the one-cell stage with only 1-2 pg of each mRNA (Figure 1b, Material and Methods), 30 times less than what has been used for the assessment of effective diffusivities in FRAP experiments^26^ and 60 times less than what is required to induce a full body axis in zebrafish^38^. In addition, we co-injected mRNA encoding membrane-targeted green fluorescent protein^44^ (memGFP) to visualize cell outlines. After injection, embryos were incubated in JF549^45^ dye solution to covalently label the HaloTag fusion protein. Subsequently, we extensively washed the embryos to remove unbound dye (Figure 1b, Material and Methods).

**Figure 1.**
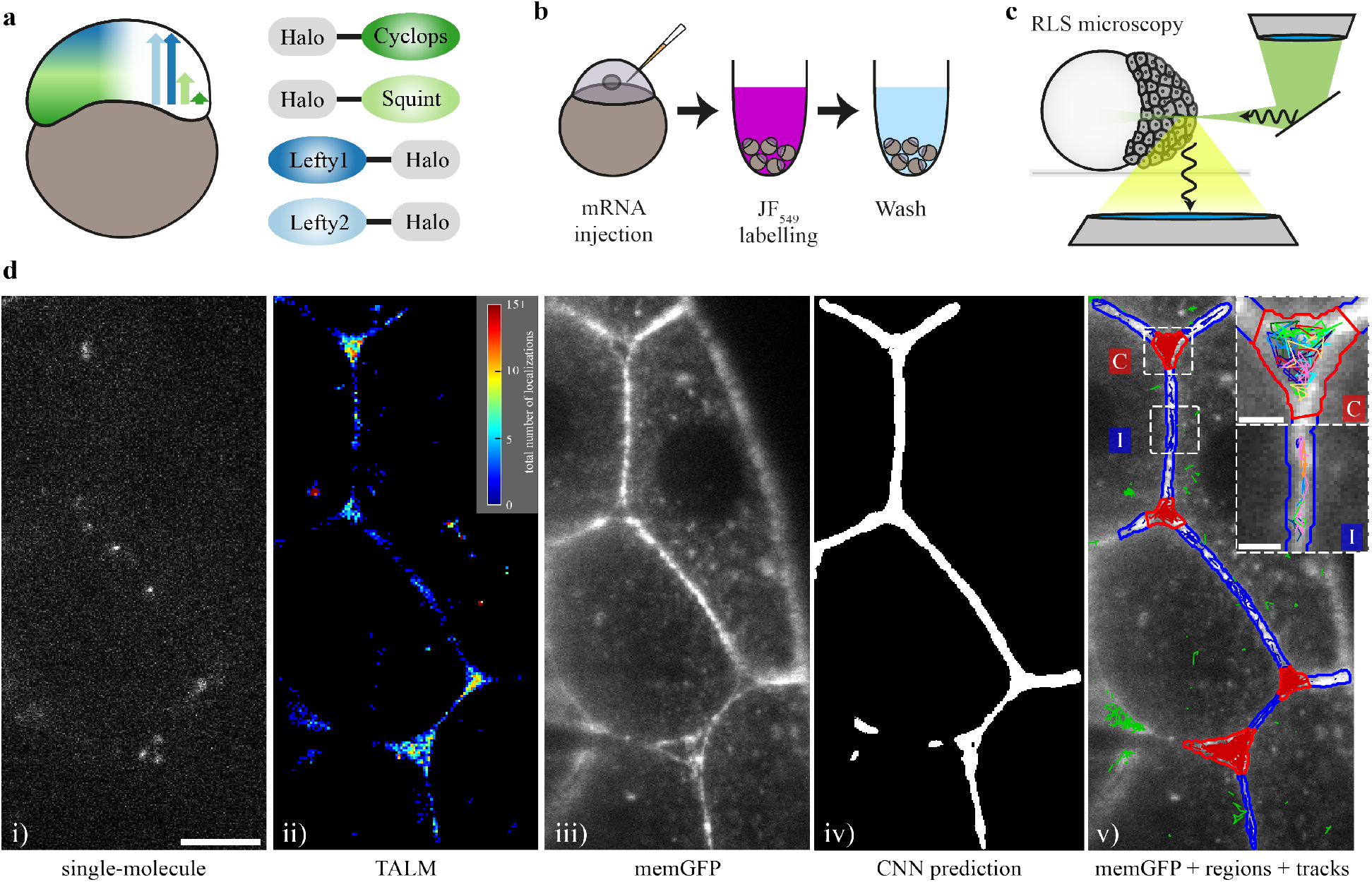
Single-molecule imaging of HaloTag-labeled Nodal and Lefty in live zebrafish embryos using RLSM. **a)** Schematic of the presumed concentration gradients of Nodal (dark and light green) and Lefty (dark and light blue) in early zebrafish embryos (left) and sketch of HaloTag fusion proteins of mature Cyclops, mature Squint, Lefty1 and Lefty2 (right). **b)** Schematic of the labeling workflow: mRNA encoding the fusion proteins was injected at the 1-cell stage. Embryos were incubated in medium containing the HaloTag JF-549 ligand to covalently label the HaloTag. Excess dye was removed in washing steps. **c)** Sketch of a zebrafish embryo imaged with a reflected light-sheet microscope. **d)** Workflow of single-molecule imaging and image segmentation: i) signal of single Lefty2-HaloTag molecules at 561 nm laser illumination for 10 ms; ii) tracking-and-localization-microscopy (TALM) image showing the total number of localizations over 1000 frames or 11.7 sec in each 2×2 pixel bin; iii) memGFP signal at 488 nm laser illumination averaged over 10 × 10 ms frames, outlining cellular membranes; iv) region of interest (ROI) mask of the extracellular space predicted by a convolutional neural net (CNN) based on the memGFP signal; v) overlay of the memGFP signal, the manually curated ROI separating cell-cell-interfaces (blue, I) and extracellular cavities (red, C); tracks assigned to interfaces (blue), cavities (red) and not assigned (green) are also shown. Scale bar: 10 µm. Insets in v): zoom of the indicated interface and cavity including an example set of tracks. Scale bar: 2 µm.

We started to image the embryos with our RLSM setup at the end of the 128-cell stage (Figure 1c) and continued the measurements up to sphere stage, shortly before gastrulation^46^. All fluorophores detected in each frame were used to track molecules in time using the software TrackIt^47^ (Material and Methods). Given the compartmentalization of the embryonic tissue into intra- and extracellular regions, we performed our tracking analysis in separate sub-regions. To automatically identify extracellular regions, we analyzed memGFP images by training a convolutional neural network (CNN)^48^ with a manually annotated data set (Figure 1d, Material and Methods). The intra- and extracellular masks classified in this manner were visually inspected and manually corrected. On average 33% of a prediction mask was truncated and subsequently 8% manually corrected. The early blastoderm comprises loosely packed cells^49^, subdividing the extracellular space into regions of close cell-cell contacts and large intercellular cavities where cell contacts are missing. We therefore further manually classified the extracellular space into interface and cavity regions and performed our single-molecule analysis separately in those regions (Figure 1d, Material and Methods).

### Nodals have similar diffusion coefficients but higher immobile fractions compared to Leftys

We first characterized the mobility of Nodals and Leftys in interfaces and cavities of the extracellular space by acquiring continuous movies of each morphogen at a rate of 85 frames per second (Supplementary Movie 1). When the single-molecule positions were integrated over all frames, the distributions of Nodals and Leftys well resembled the known localizations^26^: Cyclops was largely found in puncta, Squint in both puncta and diffusely, whereas Lefty1 and Lefty2 nearly uniformly occupied the extracellular space (Figure 2a, Supplementary Figure 1, Supplementary Movies 2-5). Interestingly, all secreted molecules were more likely to be found in cavities than in interfaces based on the ratio of localization densities as a measure of the probability to encounter a morphogen in one of the two extracellular compartments (Figure 2b), and we found an increase in the localization density ratio commensurate with the morphogens’ effective global diffusivities^26^ (Cyclops: 1.53-fold, Squint: 2.66-fold, Lefty1: 3.83-fold, Lefty2: 4.03-fold, sec-Halo: 4.20-fold).

**Figure 2.**
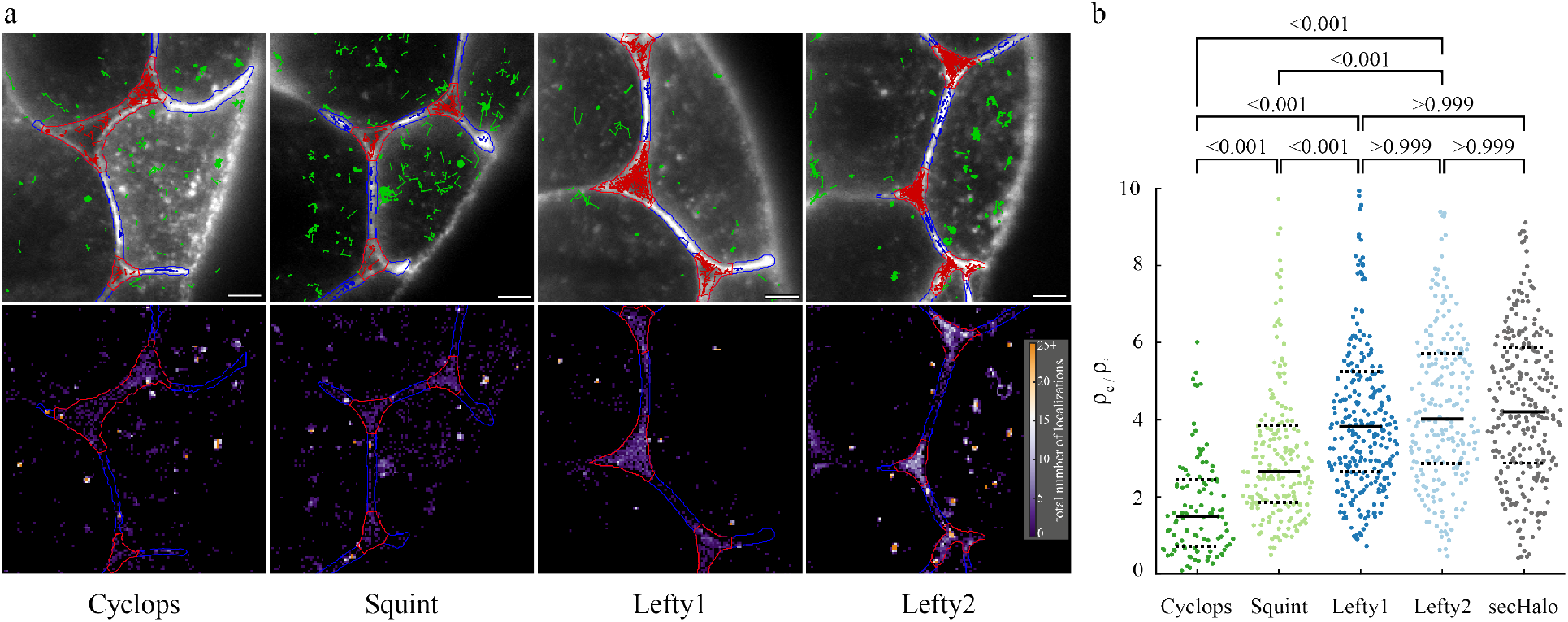
Localization of morphogens in cell-cell interfaces and extracellular cavities. **a)** Top: Overlay of the memGFP signal, the manually curated ROI separating cell-cell-interfaces (blue, I) and extracellular cavities (red, C); tracks assigned to interfaces (blue), cavities (red) and not assigned (green) are also shown. Bottom: Tracking-and-localization-microscopy (TALM) image showing the total number of localizations over 1000 frames or 11.7 sec in each 2×2 pixel bin; scale bar: 5 µm. **b)** Ratios of localization densities in cavities and interfaces calculated for each movie (for statistics see Supplementary Table 5). Values above 10 are not displayed (3.4 % of all movies). P-values were calculated using the Kruskal-Wallis-Test. Solid black lines indicate the median values, dashed black lines the 0.25 and 0.75 quantiles.

We then sorted the distances between consecutive fluorophore localizations within a track (jump distances) into histograms (Figure 3a). In both interfaces and cavities, Cyclops, and to a lesser degree Squint, showed a larger probability of short jump distances (< 0.3 µm) than Lefty1 and Lefty2, indicating reduced mobility of Nodals. In contrast, in cavities, both Nodals and Leftys exhibited a higher probability of long jump distances – and hence high mobility – compared to interfaces. We quantified the mobility of morphogens by analyzing the corresponding cumulative distribution of jump distances^50^ (Figure 3b).

**Figure 3.**
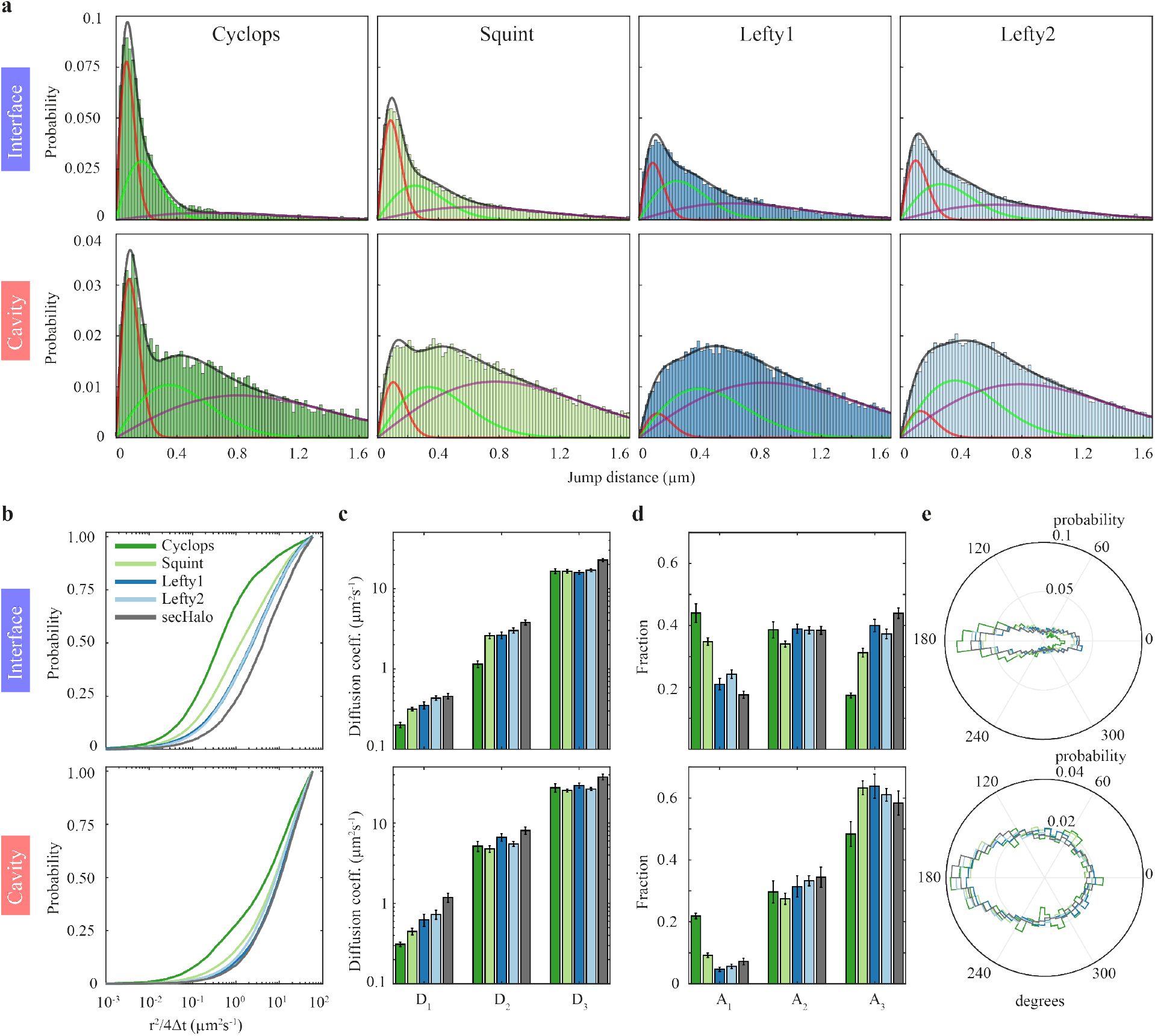
Mobility of morphogens in cell-cell interfaces and extracellular cavities. **a)** Distribution of jump distances within single-molecule tracks in interfaces and cavities for the indicated morphogen. Lines represent a three-component diffusion model (black) and the individual components (red, green, purple). **b)** Cumulative distributions of jump distances. **c)** Diffusion coefficients and **d)** fractions of the three-component diffusion model (Supplementary Table 1 and 2). Error bars denote s.d. of 500 resamplings with randomly selected 50% of the data. **e)** Distribution of angles between two consecutive track segments. For full experimental statistics see Supplementary Table 5.

A three-component diffusion model best described the data (Supplementary Figure 3), yielding the diffusion coefficients *D*_*1,2,3*_ of slow, intermediate and fast diffusion and their relative amplitudes *A*_*1,2,3*_. The slow diffusion component likely originates from immobile or slowly moving morphogens, although the uncertainty of localizing single molecules may also be a contributor. We found that in both interfaces and cavities, the diffusion coefficients of intermediate (*D*_*2,i*_ in interfaces ≈ 1.2 – 3.0 µm²s^-1^, *D*_*2,c*_ in cavities ≈ 4.8 – 6.8 µm²s^-1^) and fast (*D*_*3,i*_ ≈ 16.0 – 17.0 µm²s^-1^, *D*_*3,c*_ ≈ 25.8 – 30.0 µm²s^-1^) diffusion were largely comparable between all four morphogens, yet overall higher in cavities (Figure 3c, Supplementary Table 1 and Supplementary Table 2).

Interestingly, Cyclops showed lower diffusion coefficients in the immobile and intermediate diffusion classes, in agreement with its punctate localization pattern (Supplementary Figure 1) and potentially indicating higher confinement of this morphogen. Diffusion of the HaloTag alone (secHalo) was faster than any morphogen in both interfaces and cavities in accord with its smaller size and inert nature.

The fraction of immobile molecules in interfaces and cavities was significantly larger for Cyclops (*A*_*1,i*_ in interfaces 44%, *A*_*1,c*_ in cavities 22%) and Squint (*A*_*1,i*_ 35%, *A*_*1,c*_ 9%) than for Lefty1 (*A*_*1,i*_ 21%, *A*_*1,c*_ 5%) and Lefty2 (*A*_*1,i*_ 24%, *A*_*1,c*_ 6%) (Figure 3d), reflecting the higher probability of short jump distances for Nodals (Figure 3a). Correspondingly, while the fraction of molecules with intermediate diffusivity was similar for Nodals and Leftys, the fraction of fast-diffusing molecules in interfaces and cavities was larger for Lefty1 (*A*_*3,i*_ 39%, A_3,c_ 63%) and Lefty2 (*A*_*3,i*_ 37%, A_3,c_ 61%) than for Cyclops (*A*_*3,i*_ 17%, *A*_*3,c*_ 48%). The fraction of fast-diffusing Squint molecules was lower than that of Leftys in interfaces, but comparable to the fraction of fast-diffusing Leftys in cavities (*A*_*3,i*_ 31%, *A*_*3,c*_ 63%) (Figure 3d, Supplementary Table 1 and Supplementary Table 2). Taken together, our analysis of the diffusion data confirms that the free diffusion coefficients of Nodals and Leftys are comparable^2,15^ and suggests that the differential mobility of Nodal and Lefty reported previously^2,26,38,40^ originates from a higher retention of Nodal in an immobile state. This retention is more efficient in interfaces, where the fractions of immobile molecules are larger and diffusion is slower than in cavities.

To explore the differential diffusion properties in interfaces and cavities, we calculated the angles within a track spanned by three consecutive localizations^51^. We only considered angles where the two jumps making up the angle covered a minimum distance of 1 px (166 nm), much larger than the localization error. In interfaces, both Nodal and Lefty showed an anisotropic angle distribution with a high probability to continue or reverse the previous direction (Figure 3e). In contrast, the angle distribution was more isotropic in cavities (Figure 3e). These angle distributions probably reflect constrained diffusion along cell membranes in interfaces and free Brownian motion in cavities. The angle distributions of both interfaces and cavities exhibited a prominent contribution of reverse motion, probably reflecting partial trapping or immobilization in interfaces and free diffusion in the limited space of cavities (Supplementary Figure 4).

### Nodals bind in the extracellular space with retention times of ten to twenty seconds

To test the idea that Nodals are trapped in cell-cell interfaces, we next characterized the residence times of Cyclops and Squint in the bound state. We used time-lapse imaging, where two images are separated by dark times of different duration (Figure 4a, Material and Methods). With this illumination scheme, it is possible to increase the measurable range of binding times and to resolve several photobleaching-corrected dissociation rate constants^47,52–54^. We chose frame cycle times of 11.7 ms, 58 ms, 199 ms and 1006 ms and were thus able to observe binding events of tens of seconds along the membrane for both Cyclops and Squint (Figure 4).

**Figure 4.**
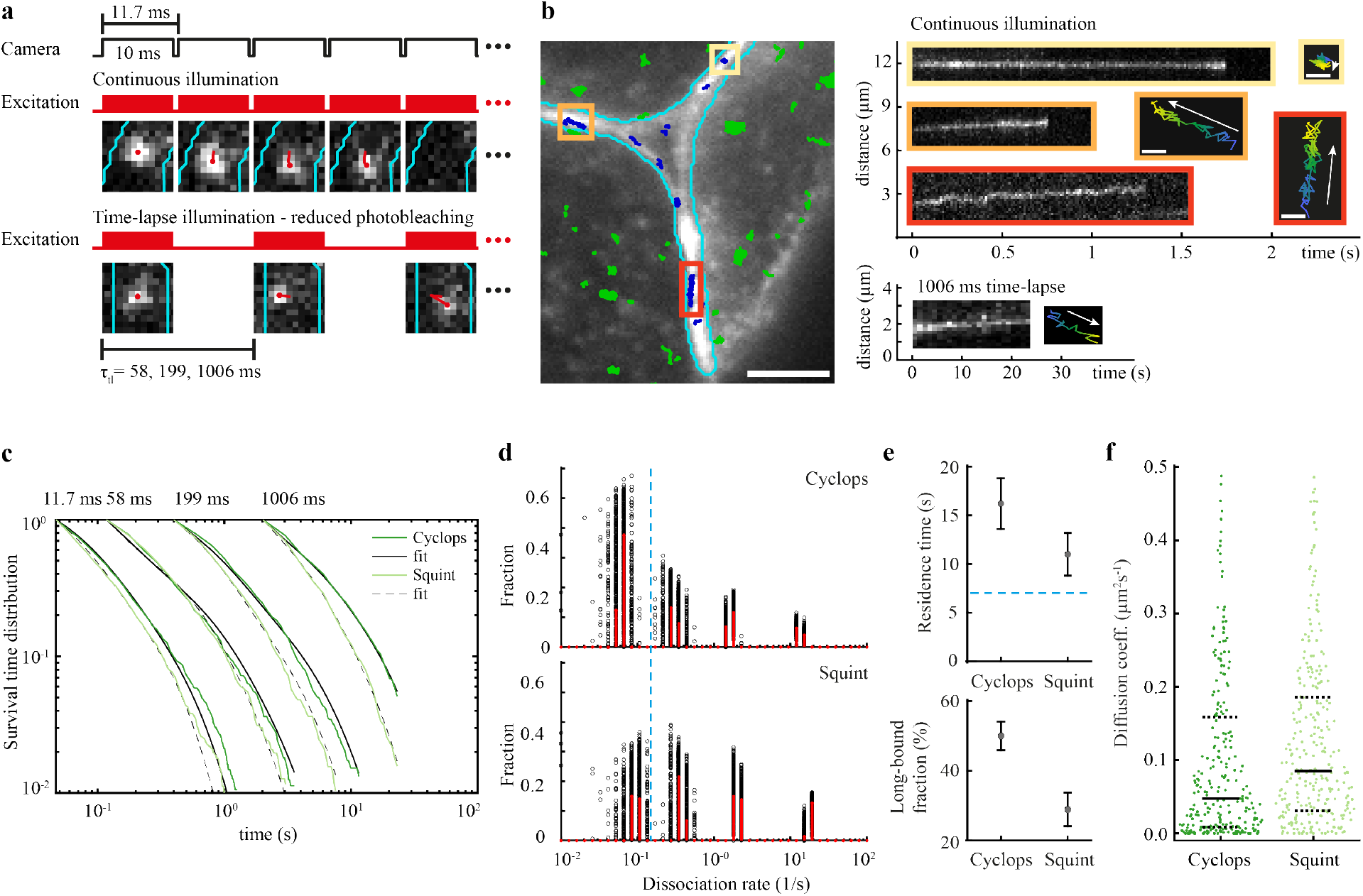
Residence times of Cyclops and Squint in the extracellular space. **a)** Overview of camera exposure and laser excitation patterns in time-lapse illumination experiments and representative images of single molecules overlaid with tracks (red) and the boundary of the extracellular region (cyan). **b)** Left panel: Overlay of the memGFP signal (white) with the boundary of the extracellular region (cyan) and tracked cyclops molecules from a movie with continuous illumination. Bound Cyclops molecules in the extracellular space shown in blue, masked, mostly intracellular molecules shown in green. Scale bar: 5 µm. Right panels: continuous illumination kymographs of the yellow, orange and red regions indicated in the left image and below a kymograph from a 1006 ms time-lapse movie as well as position plots of the tracks color-coded according to start (blue) and end (yellow) times. Scale bar: 0.3 µm. **c)** Survival-time distributions of bound morphogens in the time-lapse conditions indicated on top (dark and light green lines) and survival time functions obtained by GRID for Cyclops (solid black lines) and Squint (dashed black lines). **d)** State spectra of dissociation rates of Cyclops and Squint obtained by GRID using all data (red bars) and 500 resampling runs with randomly selected 80% of data (black data points) as an error estimation of the spectra. The dashed blue line indicates the boundary for long-bound molecules of 0.15 s^-1^. **e)** Residence times of Cyclops and Squint (top panel) and the corresponding fraction (bottom panel) extracted from the slowest dissociation rate cluster of the state spectra. Error bars denote s.d. of the resampled spectra in d). The dashed blue line indicates the boundary for long-bound molecules of 7 s. **f)** Diffusion coefficients obtained from fitting the first 10 points of a mean-squared displacement plot of bound Cyclops and Squint molecules recorded for at least 20 frames (n_tracks,cyc_ = 283, n_tracks,squ_ = 337). Values above 0.5 µm²s^-1^ were discarded (11%). Solid black lines indicate the median values, dashed black lines the 0.25 and 0.75 quantiles. For full experimental statistics see Supplementary Table 6.

We identified bound molecules in interfaces and cavities using a small tracking radius in combination with a minimum number of survived frames in the nearest neighbor algorithm^47^ (Material and Methods). We then collected the durations of binding events for each time-lapse condition in survival-time distributions (Figure 4c, Material and Methods). The distributions extended to longer durations for Cyclops than for Squint, indicating longer binding times for Cyclops. For the longest time-lapse condition, where photobleaching is not limiting, few binding events survived throughout the whole acquisition time (5% for Cyclops, 1.4% for Squint). Thus our analysis will slightly underestimate the binding times. Lefty1 and Lefty2 exhibited much-reduced occurrences and durations of binding events in movies of 11.7 ms frame cycle time, comparable to those of the HaloTag alone (Supplementary Figure 6b). Together with the low bound fractions obtained from the diffusion analysis, this indicates that binding in the extracellular space has a minor influence to the overall diffusion properties of Leftys, and we therefore refrained from quantifying their binding times. For Cyclops and Squint, we analyzed the survival-time distributions with GRID, which can reveal spectra of dissociation rates from fluorescence survival-time distributions by solving the inverse Laplace transformation^52^. We obtained four dissociation rate clusters for both Cyclops and Squint (Figure 4d), from which the inversely correlated binding times can be calculated. The longest binding time, corresponding to the slowest dissociation rate cluster, was 16.2 ± 2.6 s (mean ± s.d. of resampled spectrum), comprising 50.0 ± 4.1 % (mean ± s.d. of resampled spectrum) of bound molecules for Cyclops. For Squint, we found shorter binding times of 11.0 ± 2.2 s comprising 28.9 ± 4.8 % of bound molecules (Figure 4e), consistent with the larger effective diffusion coefficient, the less punctate distribution compared to Cyclops^26^ (Supplementary Figure 1), and in remarkable agreement with the previous dissociation rate predictions of 18 s for Cyclops and 4 s for Squint^26^.

Some of the Cyclops and Squint molecules that we identified as bound showed slow diffusive motion along the membrane (Figure 4b, Supplementary Movies 6 and 7). This observation is in agreement with a fraction of slowly diffusing morphogens obtained in the analysis of molecular jump distances (Figure 3c,d). Such motion might correspond to the diffusion of morphogen-receptor complexes within the membrane. To test this idea, we quantified the diffusion coefficients of bound morphogens by analyzing the mean-squared displacement (MSD) of bound tracks for different time intervals (Supplementary Figure 5, Supplementary Movies 6 and 7, Material and Methods). The majority of diffusion coefficients of both Cyclops and Squint was below 0.5 µm^2^s^-1^ (Figure 4f), indeed similar to previous quantifications of receptor diffusion in membranes^55,56^. Bound Squint molecules exhibited a higher tendency to diffuse along the membrane than bound Cyclops molecules, which were more often confined to a small area, again consistent with the larger effective diffusion coefficient and less punctate distribution compared to Cyclops^26^ (Supplementary Figure 1).

### Overexpression of oep increases the fraction of immobile Squint molecules

What could be the molecular interaction that leads to the higher retention times of Nodals compared to Leftys? Nodals are well known to bind to the EGF-CFC co-receptor Oep^37,57^, which is essential for Nodal signaling^58^. Furthermore, the signaling range of Squint was shown to be extended in the absence of *oep*^41,42^, but a direct effect of Oep on Nodal dispersal has not yet been directly demonstrated. To test whether immobile Squint in our experiments was due to binding to Oep, we co-injected 0.3 pg, 3 pg and 30 pg of Oep-encoding mRNA together with the Squint-HaloTag construct and compared the diffusion properties of Squint in conditions of *oep* overexpression with those of endogenous Oep levels.

We found that the immobile fraction of Squint increased in interfaces and cavities, from 35% to 50% and from 9% to 25% respectively (Figure 5b and Supplementary Figure 7a), while the diffusion coefficients remained unaffected (Figure 5a). Thus, our data show on a single-molecule level that Oep can directly hinder the diffusion of Nodal by transiently trapping the morphogen on the membrane.

**Figure 5.**
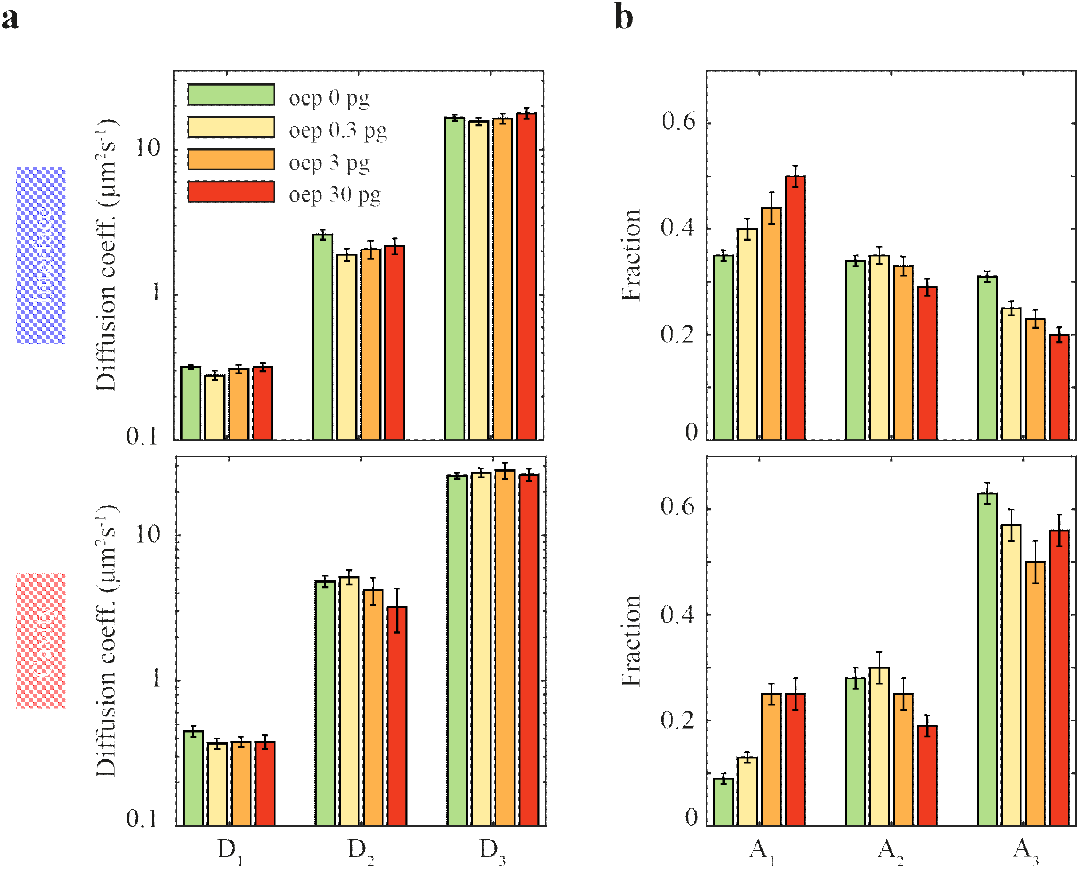
Mobility of Squint decreases with overexpression of *oep*. **a)** Diffusion coefficients and **b)** fractions of the three-component diffusion model in interfaces and cavities for Squint at different amounts of mRNA encoding *Oep*. Error bars denote s.d. of 500 resamplings with randomly selected 50% of the data. For full experimental statistics see Supplementary Table 5.

### Clarifying origins of differential morphogen localization using agent-based modeling

Interestingly, we found a higher fraction of immobile molecules combined with slower diffusion in cell-cell interfaces compared to extracellular cavities. These findings would intuitively suggest that morphogens should accumulate in interfaces, not cavities. However – surprisingly, and in contrast to intuition – we found that all secreted molecules were more likely to be found in extracellular cavities rather than in cell-cell interfaces. Mathematical modeling can help reveal the origins of non-intuitive behaviors in biological systems^38,40,59^. We therefore devised a minimal model of single-molecule dispersal in order to test whether geometric constraints and binding might suffice to explain the cavity enrichment, or whether more complicated molecular mechanisms such as restricted entry control into interfaces have to be invoked.

To simulate single-molecule dispersal, we chose an agent-based model in a realistic zebrafish blastoderm geometry that directly relates to our experimental observations. We used an experimentally determined binary mask of extracellular space as two-dimensional grid (Figure 6a). Single morphogens were simulated as “drunken sailors”^2^ performing a random walk in the extracellular space. To simulate immobile and freely diffusing single molecules, we used jump sizes at each simulation step based on the measured diffusion coefficients for bound (0.5 µm^2^s^-1^) and free (30 µm^2^s^-1^) states (see Material and Methods). A single molecule became bound when it detected a receptor in proximity (≤ 20 nm). The simulated tracks for Nodals (Figure 6a, blue track) and Leftys (Figure 6a, red track) closely resembled the experimental observations. In particular, our simulation was able to recapitulate higher bound fractions in interfaces compared to cavities, differential angle distributions in both compartments as well as higher localization density in cavities compared to interfaces (Figure 6 b-c, Supplementary Figure 8 a-f).

**Figure 6.**
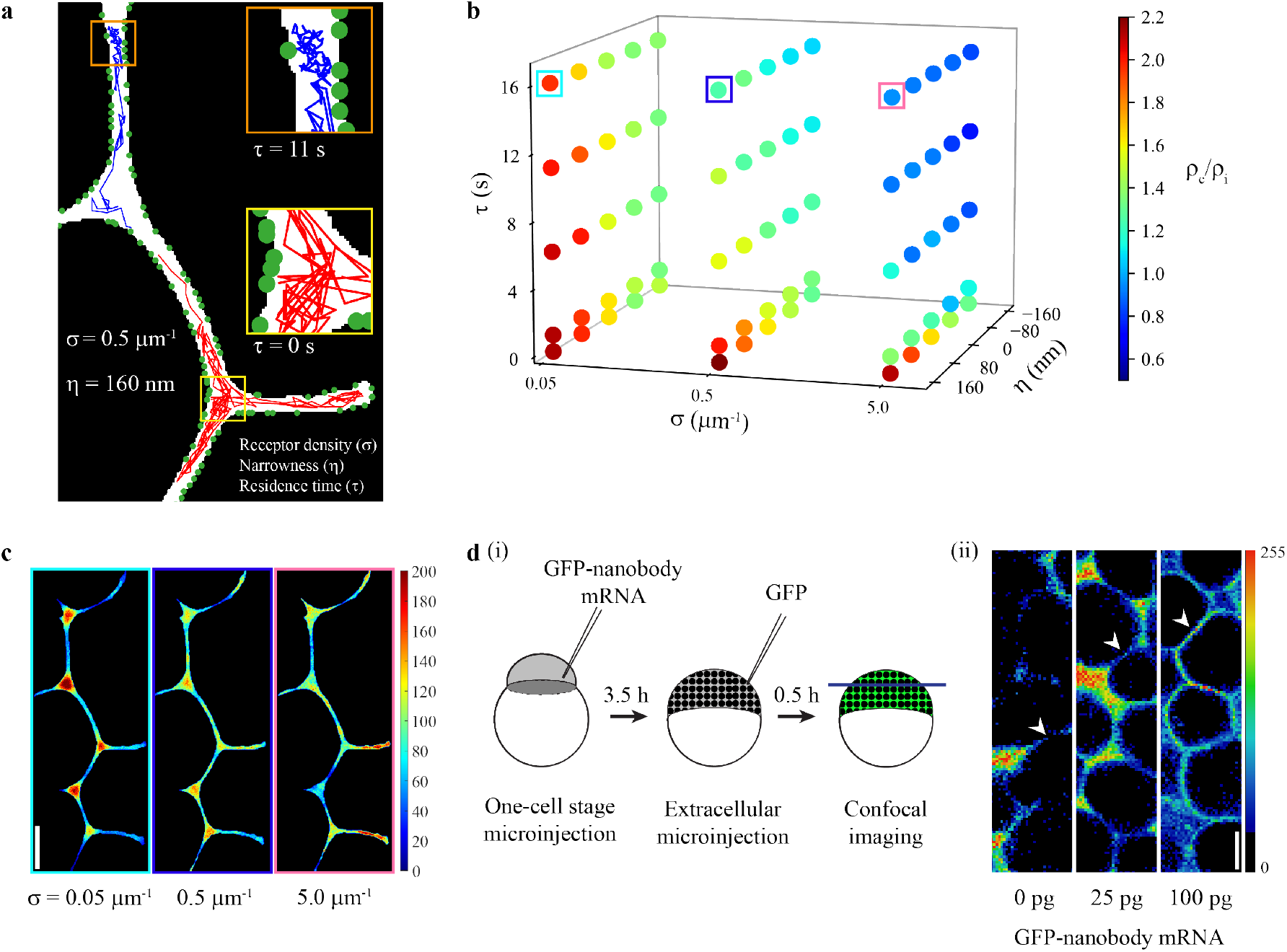
An agent-based model reveals key parameters affecting morphogen behavior in the extracellular space. **a)** Illustration of the model with a section of the two-dimensional grid showing simulated tracks (200 steps or 2 s) for two morphogens with residence times *τ* of 11 s (blue track) and 0 s (red track), respectively. Green circles indicate receptors. The receptor density σ is given as the number of receptors per µm of membrane length. The width of the extracellular region is reduced by the narrowness factor *η*, Morphogens are allowed to move in the extracellular space (white). A morphogen becomes ‘immobile’ or bound upon detecting a receptor in proximity and performs smaller jumps corresponding to the bound diffusion coefficient. **b)** Three-dimensional scatter plot of simulated parameter combinations showing corresponding localization density ratios (ρ_c_/ρ_i_). Five residence times *τ* (0 s, 1 s, 6 s, 11 s, and 16 s), three receptor densities σ (0.05 µm^-1^, 0.5 µm^-1^, and 5.0 µm^-1^) and five narrowness conditions *η* (−160 nm, -80 nm, 0 nm, 80 nm, and 160 nm) were simulated. The average width of the extracellular space was approximately 1 µm. Negative *η* values indicate that the space was widened. **c)** Localization density plots of simulated morphogens with increasing receptor densities σ (0.05 µm^-1^, 0.5 µm^-1^, and 5.0 µm^-1^; *τ* = 16 s, *η* = 160 nm). Frame colors correspond to colored boxes in b). **d)** (i) Schematic depicting the experimental procedure to acquire confocal images of GFP localization in zebrafish embryos. (ii) Representative images of regions of interest used for GFP signal measurements. Cell-cell interface regions are indicated by arrowheads. Scale bars in c and d (ii) are 10 µm.

Next, we systematically varied key parameters that might modulate morphogen localization in the extracellular space. A parameter screen revealed that receptor density (σ), residence time (*τ*), and interface narrowness (*η*) are important determinants of extracellular molecule localization (Figure 6b). As expected, strong binding (σ > 0.5 µm^-1^ and *τ* > 6 s) yielded localization in cell-cell interfaces (Figure 6 b,c and Supplementary Figure 8a). Surprisingly, for low binding (σ < 0.5 µm^-1^ and *τ* < 6 s) extracellular molecules tended to localize in cavities. This tendency was augmented by decreasing the width of the extracellular space, suggesting an important role of tissue geometry for morphogen localization (Figure 6b and Supplementary Figure 8 b). We verified the accumulation in cavities for multiple additional experimentally derived grids (Supplementary Figure 8 c). The narrow width of interfaces concentrated extracellular molecules to cell surfaces, thereby increasing the probability of interactions with receptors. This behavior resulted in a higher bound fraction in interfaces compared to cavities despite similar receptor spacing in both compartments (Figure 3d, Supplementary Figure 8 d,e).

Our simulations showed that the surprising enrichment of extracellular molecules in cavities can be explained purely by geometric constraints, and predicted that they could be pushed out of this compartment into interfaces by increasing receptor density homogeneously in the tissue (Figure 6c). To test this prediction, we measured the distribution of extracellular GFP in zebrafish embryos with different levels of membrane-tethered GFP-binding nanobodies^60^. Similar to our TALM findings (Figure 1d and Figure 2), secreted GFP was mainly distributed in cavities (ρ_c_/ρ_i_ > 2) in case of no or low amount (25 pg) of injected nanobody mRNA. With high nanobody expression (100 pg), GFP signal became more enriched in cell-cell interfaces (ρ_c_/ρ_i_ < 1) (Figure 6d and Supplementary Figure 8g), in accordance with our simulations. Our results suggest that geometric constraints, such as interface narrowness, bias morphogens to preferably localize in extracellular cavities. Strong binding to receptors can overcome this bias to increase morphogen localization in cell-cell interfaces.

Together, our single-molecule measurements and simulations provide strong support for the hindered diffusion model, in which differences in effective diffusivities between Nodals and Leftys are an emergent property arising from differential binding of morphogens in a compartmentalized extracellular environment.

## Discussion

Multiple mechanisms have been proposed to underlie the dispersal of morphogens for developmental patterning, from simple extracellular diffusion to repeated secretion and cellular uptake, filopodia-based distribution, and signal relay_^2,9,14,15,31,61–68^_. In particular, the hindered diffusion model postulating free diffusion intermitted by transient binding on cell surfaces has gained popularity^7^. Dispersal models have been inferred from observations of large averaged morphogen ensembles and bulk mobility measurements using techniques such as FCS or FRAP. However, the resulting data has to be carefully interpreted because bulk measurements may only provide indirect evidence for transport mechanisms^39,69–71^. To directly determine the mechanisms of transport, single-molecule experiments are necessary, but these measurements have so far not been performed for any morphogen in a living embryo.

We performed single-molecule measurements of individual Nodal and Lefty morphogens in developing zebrafish embryos. Our results suggest that morphogens undergo free and fast Brownian motion in extracellular cavities, whereas diffusion is constrained and slower in cell-cell interfaces. The coefficients of free diffusion were similar for Nodals and Leftys. In contrast, Nodals, and in particular the ultra-short-range morphogen Cyclops, exhibited a larger fraction of molecules bound to the cell membrane than Leftys. Our direct single-molecule observations are consistent with previous inference from indirect bulk measurement techniques such as FCS and FRAP^2,15,26,39^, and here we show both diffusion and reversible morphogen binding with single-molecule resolution in strong support of the influential hindered diffusion model.

Surprisingly, we observed an unexpected accumulation of morphogens in extracellular cavities, although binding interactions were more pronounced in cell-cell interfaces than in cavities. Using agent-based simulations of morphogen transport, we found that the architecture of the extracellular space with large cavities and narrow cell-cell interfaces favors heterogeneous distribution of morphogens and their accumulation in cavities. Our simulations predicted that binding to receptors would counteract this effect, and we validated this idea by measuring the distribution of extracellular GFP in zebrafish embryos with different levels of membrane-tethered artificial GFP-binding receptors^60^. In addition, stronger or more frequent binding, implemented by longer residence times or higher receptor densities, respectively, retained morphogens in cell-cell interfaces. Furthermore, the narrow width of interfaces contributed to enhanced binding by concentrating morphogens to cell surfaces. The influence of tissue architecture on effective diffusion coefficients has been discussed in previous studies^2,15,39^: Numerical simulations and experiments using FCS and FRAP with secreted GFP inferred a reduction of effective diffusion compared to free diffusion by a factor of approximately two-fold^14,26,39^, rationalizing the idea that secreted molecules have to bypass other cells compared to diffusion in free space. Our new direct single-molecule measurements and simulations suggest a molecular mechanism of long-range morphogen spreading in a tortuous environment, where diffusion predominantly occurs in a network of extracellular cavities. This mechanism differs from the morphogen dispersal mode recently described in a two-dimensional human embryonic stem cell culture system, in which – unlike in the three-dimensional embryo context – Nodal molecules presumably cannot be retained in extracellular cavities and are instead lost into the culturing medium^66^.

We observed that binding differed between the Nodals Cyclops and Squint. First, the bound fractions where higher for Cyclops than for Squint, and binding times of Cyclops were on average 5 s longer. This contrasts with measurements showing that Squint binds to the Type II receptor Acvr2b-a with higher affinity than Cyclops^15^. Second, we observed that bound Squint molecules frequently exhibited slow diffusion along the membrane, while bound Cyclops molecules were mostly localized within a small area and effectively immobilized. Our single-molecule measurements are oblivious to the molecular identity of the morphogen binding partners and therefore represent a neutral description of overall cell surface binding. The differing mobilities of bound Cyclops and Squint might reflect differences in how both members of Nodal bind to components of the extracellular matrix. There are numerous potential extracellular binding partners, for example the EGF-CFC co-receptor Oep or other immobilized diffusion regulators such as heparan sulfate proteoglycans^7,8,18–22,60,72^. The different degrees of hindered diffusion that we observed – shorter residence times and lower fraction of bound Squint compared to Cyclops – are likely to underlie the different ranges of Nodal gradients (short-to-mid range for Squint and ultra-short range for Cyclops)^26,73–76^.

Interestingly, while we also observed labeled Nodals and Leftys in the cytoplasm (excluded in the present analysis) in addition to their extracellular localization, we only rarely (in approximately 3 out of 100 movies) observed events where single molecules clearly passed the membrane and entered the cytoplasm. This indicates that internalization of Nodals/Leftys is a rare event, as opposed to the prolonged binding of Nodals on the cell surface, which provides a possible explanation for the hour-long half-lives of Nodals/Leftys in living zebrafish embryos^26,42,77^. Rare internalization also contrasts with the transcytosis mechanism described for the TGF-β superfamily ligand Dpp in *Drosophila*, in which repeated round of exocytosis and endocytosis lead to morphogen dispersal^63^. The differences in the dispersal mechanisms might be explained by the different time scales required for patterning of the zebrafish embryo (hours) and the *Drosophila* wing disc (days)^2^.

In summary, we propose that Nodal and Lefty spreading follows a compartmentalized hindered diffusion model, in which cell-cell interfaces provide a confined, obstructive environment with restricted diffusion in particular for Nodal, whereas long-distance spreading of morphogens occurs within cavities between cells.

## Material and Methods

### Generation of constructs

The designs for the HaloTag-tagged zebrafish Nodals Squint and Cyclops, Lefty1 and Lefty2 as well as secreted HaloTag were based on previously published GFP fusion constructs^26^.

To generate pCS2-2xHA-HALO-3xGs, HaloTag was isolated using primers containing a tandem HA-Tag and primers containing a triple GS-linker, which was then cloned into the BamHI and XbaI sites of the pCS2 backbone^43^.

HaloTag was then amplified from pCS2-2xHA-HALO-3xGS, and fusion constructs for Nodals and Leftys were generated using splicing-by-overlap-extension PCR^78–80^ using the pCS2 backbone. The following primers were used:

For pCS2-2xHA-HALO-3xGs:

GATCGGATCCATGTACCCATACGATGTTCCAGATTACGCTGGATATCCATATGATGTTCCAGATTATGCTCGAGGAGCAGAAATCGGTACTGGCTT, GATCATCTAGAGATCGAGGCGCGCCGATCGATTAATTAAGCTTCCGGAGCCAGAACCTGAGCCGGAAATCTCGAGCG

For pCS2-Squint-HaloTag:

GATGGATCCACCGGTACCACCTCGACCTCCATCACGGCC, GGCTCGAGAGGCCTTGAATTCTCAGTGGCAGCCGCATTCTGC, CTCGAGATTTCCGGCGGATCCGCAGCAGCAG, CTGCTGCTGCGGATCCGCCGGAAATCTCGAG, GATCCACCGGTACCACCGGAGCAGAAATCGGTAC, GTACCGATTTCTGCTCCGGTGGTACCGGTGGATC

For pCS2-Cyclops-HaloTag:

GCAGGATCCCATCGATGCCACCATGCACGCGCTCGGAGTCGC, GGCTCGAGAGGCCTTGAATTCTCACAGGCATCCGCACTCCTC, GCCGCCGGGGGCCAGGAGCAGAAATCGGTAC, GTACCGATTTCTGCTCCTGGCCCCCGGCGGC, CTCGAGATTTCCGGCCCTGTCAGGAGCCCAG, CTGGGCTCCTGACAGGGCCGGAAATCTCGAG

For pCS2-Lefty1-HaloTag:

GCAGGATCCCATCGATGCCACCATGACTTCAGTCCGCGCCG, CTATAGTTCTAGAGGCTCGAGTCAGCCGGAAATCTCGAG, GATCCACCGGTCGCCACCGGAGCAGAAATCGGTAC, GTACCGATTTCTGCTCCGGTGGCGACCGGTGGATC

For pCS2-Lefty2-HaloTag:

GCAGGATCCCATCGATGCCACCATGGCTCTGTTCATCCAGC, CTATAGTTCTAGAGGCTCGAGTCAGCCGGAAATCTCGAG, CCCTCCAGTCCTGGGCGGAGCAGAAATCGGTAC, GTACCGATTTCTGCTCCGCCCAGGACTGGAGGG

To generate pCS2-secreted-HaloTag, HaloTag was amplified with the primers GATCCACCGGTACCACCGGAGCAGAAATCGGTAC and CTATAGTTCTAGAGGCTCGAGTCAGCCGGAAATCTCGAG and cloned by restriction digest using AgeI and XbaI.

### mRNA synthesis

For capped mRNA synthesis, pCS2-Cyclops-HaloTag, pCS2-Squint-HaloTag, pCS2-Lefty1-HaloTag, pCS2-Lefty2-HaloTag, pCS2-secreted-HaloTag, pCS2-Squint-GFP^26^ and pCS2-mem-GFP^44^ were linearized with NotI, and a mMESSAGE mMACHINE SP6 Kit (Invitrogen) was used for *in vitro* transcription according to the manufacturer’s recommendations. To generate mRNA encoding Oep, pCDNA3-oep-FLAG was linearized with NotI and transcribed using an mMESSAGE mMACHINE T7 Kit (Invitrogen) according to the manufacturer’s recommendations.

### Zebrafish husbandry

Wild Indian Karyotype (WIK) and TE zebrafish were maintained in accordance with the guidelines of the State of Baden-Württemberg (Germany) and approved by the Regierungspräsidium Tübingen (35/9185.46-5, 35/9185.81-5).

### Confocal fluorescence microscopy

TE embryos were injected at the one-cell stage with 50 pg of mRNA encoding secreted-HaloTag, Squint-HaloTag or Cyclops-HaloTag or 60 pg of mRNA encoding Lefty1-HaloTag or Lefty2-HaloTag. Embryos were proteolytically dechorionated using 10 mg Pronase (Sigma Aldrich) in 10 ml Danieau’s medium and washed with Danieau’s medium to remove the Pronase. The embryos were then incubated in a 1:5,000 dilution of TMR HaloTag Ligand (5 mM; Promega) in Danieau’s medium. After 30-60 min at 28°C, they were rinsed with embryo medium twice and mounted in a glass bottom dish using 1.5% low-melting agarose. Imaging was performed with an LSM 780 NLO (ZEISS) system using an LD LCI Plan-Apochromat 25*×*/0.8 Imm Korr DIC objective to acquire animal views at a depth of approximately 35 µm into the tissue.

### Sample preparation for single-molecule imaging

Embryos were dechorionated directly after fertilization using 10 mg Pronase (Sigma Aldrich) in 10 ml Danieau’s medium and washed with Danieau’s medium to remove the Pronase. To express morphogen constructs, embryos were injected at the 1-cell stage with 1 pg of mRNA encoding Squint-HaloTag, Lefty1-HaloTag, Lefty2-HaloTag, secreted HaloTag or 2 pg of mRNA encoding Cyclops-HaloTag together with 10 pg of mRNA encoding memGFP into the animal pole. The diameter of the injection mix droplet was measured with a stereo microscope (Olympus SZX2-ZB10) equipped with a camera (CAM-SC50) and the Cellsens imaging software, and set to 124 µm, corresponding to a droplet volume of 1 nl, by adjusting the injection duration. For Oep experiments, 1 pg of mRNA encoding HaloTag-Squint was injected together with 0.3 pg, 3 pg or 30 pg of mRNA encoding Oep and 10 pg of mRNA encoding memGFP.

To label the HaloTag, embryos at the 2-cell stage were placed into two separate glass tubes, and most of the embryo buffer was removed such that embryos were just covered sufficiently. 1 ml of 5 nM HaloTag-JF549^45^ dye solution was then added into one of the tubes and 10 nM HaloTag-JF549 dye solution into the other tube and incubated for 30 min. After staining, zebrafish were washed with Danieau’s buffer, followed by two additional washing steps each after 15 min.

Embryos were incubated at room temperature (22°C). The transition from the 64-cell to the 128-cell stage up to the 128-cell stage, and synchronously developing embryos were visually identified. 6-8 embryos at the 128-cell stage stained with 10 nM dye solution were then mounted onto the microscope by placing them into a glass bottom dish (Delta T, Bioptechs, Butler, PA). If the density of visible fluorescent molecules was too high, the mounted embryos were exchanged with embryos stained with 5 nM dye solution. On the microscope, embryos grew further at room temperature. Transitions between embryo stages were counted by identifying cytoplasmic divisions (cytokinesis events), when the GFP-stained cell membrane grows inward until cell division. Fluorescence imaging was stopped when embryos reached the sphere stage.

### Reflected light-sheet microscopy (RLSM)

Single-molecule imaging of live zebrafish embryos was carried out using a custom-built reflected light-sheet microscope^54^ with modifications to image live zebrafish embryos^43^. The microscope was built around a commercial Nikon TI microscope body equipped with a water-immersion objective (60× 1.20 NA Plan Apo VC W, NIKON), a dichroic mirror (F73– 866/F58–533, AHF), an emission filter (F72– 866/F57–532, AHF), a notch filter (F40–072/F40– 513, AHF) and an EM-CCD Camera (iXon Ultra DU 897U, Andor). Fluorescence light was post-magnified by a factor of 1.5x before reaching the camera chip, which resulted in a pixel size of 166 nm. The microscope was controlled using the NIS Elements software (Nikon) and a NIDAQ data acquisition card (National Instruments).

Reflected light-sheet illumination was achieved using a custom-built tower mounted above the sample dish. AOTF (AOTFnC-400.650-TN, AA Optoelectronics) controlled laser light of 488 nm (IBEAM-SMART-488-S-HP, 200 mW, Toptica) and 561 nm (Jive 300 mW, Cobolt) was coupled into the tower via a single-mode fiber, where it was focused by a cylindrical lens into the back-focal plane of a water-dipping objective (40× 0.8 NA HCX Apo L W, Leica) and subsequently reflected by the chip of an AFM cantilever (custom-coated cantilever based on model: HYDRA2R-100N-TL-20 but with both sides coated in 40 nm Al). The resulting light-sheet had a thickness of approximately 3 µm. The laser power of the light-sheet was 40 mW for the 561 nm laser and 4 mW for the 488 nm laser.

Compared to previous measurements with the mEos2 label^43^, HaloTag together with JF549 allowed for faster frame cycle times and longer tracks. Movies of morphogens were recorded with 10 ms exposure time and a total frame cycle time of 11.7 ms. The illumination time was set to match the exposure time.

For continuous movies, a sequence starting with 10 frames with 488 nm laser illumination showing the membrane-GFP signal was recorded, followed by 1000 frames of 561 nm laser illumination to image single morphogen molecules and finalized by another 10 frames of 488 nm illumination.

For time-lapse microscopy movies of Squint-HaloTag and Cyclops-HaloTag, frames were illuminated for one frame with the 561 nm laser followed by one frame with 488nm laser illumination. Dark times of different lengths were introduced between illuminated frames, resulting in frame acquisitions every 58 ms, 199 ms or 1006 ms (time-lapses) for which a total number of 200, 59 and 24 illuminated frames were recorded, respectively.

### Splitting movies into reference channel and single-molecule channel

Movies containing frames with memGFP signal as well as frames with signal of Halo-tagged morphogens were separated using TrackIts movie splitter. Two separate movies were obtained, one containing only the memGFP signal and one containing only single-molecule signal. Movies were discarded if considerable drift due to embryo movement was evident in order to guarantee a correct classification of tracks to their region classes (cavity, interface).

### Segmentation of extracellular regions using a CNN

A U-Net^81^ CNN was trained to segment extracellular regions of developing zebrafish embryos based on the memGFP intensity using ZeroCostDL4Mic^48^, a state-of-the-art image segmentation platform. Target images for training were created by first using TrackIts sub-region drawing tool to manually draw outlines around the extracellular space of 452 images.

The “Average frames” function was used to average all frames of a memGFP movie, and the “Gaussian filter” function with a kernel size of 1 px was applied to smoothen the averaged image. A custom Matlab script was then used to create a U-Net compatible 8-bit .tif file containing the training masks.

Training was performed in the cloud using the Google Colaboratory platform provided by DL4Mic. The model was trained over 200 epochs on 90% of the training images while 10% of images were used for validation. This resulted in a final training loss of 0.163. The trained model was downloaded and integrated into a custom Python program to create regions of interest compatible with our single-molecule tracking software TrackIt^47^. All frames of the memGFP movies in a folder were averaged and padded with zeros to match the U-Net network requirements. The images were then segmented with the trained model, and binary masks were created by applying a user-defined intensity threshold between 0-255, which was set to 240. Polygonal regions of interest with a minimum size of 150 px were then saved in a TrackIt compatible .roi file.

Once loaded into TrackIt, the segmentation results were visually quality-controlled for each movie. Parts where the memGFP signal was blurry (e.g. when lying far away from the edge of the zebrafish), or parts where the laser light was blocked or absorbed, were either adjusted or cut-off manually.

A second region containing extracellular cavities formed by multiple cells was added by manually selecting parts of the regions that had been segmented by the CNN (see for example Figure 1d).

### Tracking of single-molecule microscopy data for mobility analysis of morphogens

Single-molecule microscopy data of Halo-tagged Cyclops, Squint, Lefty1 & Lefty2 and secreted-HaloTag were analyzed with TrackIt^47^. A threshold factor of 1.5 was used to detect single molecule events. The nearest neighbour algorithm was used with a tracking radius of 10 px (=1.66 µm) to link single-molecule detections into tracks, and 1 gap frame was allowed to bridge detection gaps if a molecule was already detected for at least 2 consecutive frames. TrackIts “Delete tracks touching borders” option was used, which means that tracks were assigned to regions if they lied completely inside the region of interest while tracks crossing region borders were discarded and treated as non-linked detections.

### v*Distribution of jump distances and diffusion analysis*

For diffusion analysis, TrackIt’s data analysis tool was used to fit the cumulative distribution of jump distances with a three-component Brownian diffusion model. The total number of bins of the cumulative jump distance histogram was set to 1660 corresponding to a bin size of 1 nm. To prevent an overrepresentation of bound molecules, a maximum of 10 jumps was considered per track. Jumps over gap frames were not considered. The errors of diffusion coefficients *D*_*1,2,3*_ and fractions *A*_*1,2,3*_ were estimated by repeating the analysis 500 times using random samples of 50% of the jump distances, and the standard deviation of the resulting diffusion coefficients and fractions were calculated.

To visualize the diffusion analysis results, the probability distribution p(r), as obtained from the fit results, was plotted together with the histogram of jump distances using

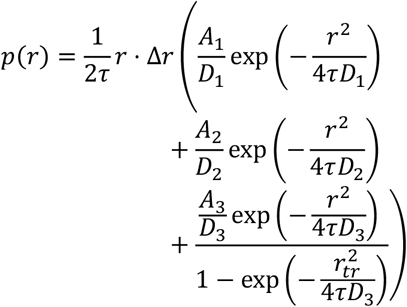

where *r* is the jump distance, Δ*r* is the bin width of the jump-distance histogram (here 27.7 nm), *τ* is the frame cycle time and *D*_*i*_ and *A*_*i*_ are the diffusion coefficients and fractions resulting from the cumulative jump distance distribution fit. The last term is normalized by 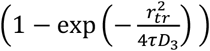, with *r* representing the tracking radius, to account for the cut-off due to the lower and upper limit of jump distances.

### Mean squared displacement (MSD) analysis

For MSD analysis tracks with a minimum duration of 20 frames were considered. The first 10 points of the MSD curves were fitted with the linear function MSD = 4 · *D* · *τ* + *c*, where *D* is the diffusion coefficient and *c* is a constant to account for the localization error.

### Spot density ratio

The density of single-molecule detections was calculated for each movie and region class separately by dividing the number of detections in each of the regions by its number of pixels. The ratio between the spot density in the cavity region and the spot density in the interface region was then calculated for each movie. Movies containing no detections in one of the regions or movies with only one region class were discarded. P-Values were calculated using the non-parametric Kruskal-Wallis-Test.

### Analysis of time-lapse microscopy data

Tracking settings were optimized for the nearest-neighbour algorithm to track only bound molecules. TrackIt’s automatic tracking radii prediction tool was used to ensure equal tracking-loss probabilities due to tracking errors and photobleaching across all time-lapse conditions. The resulting tracking radii for a loss probability of 0.005 were: 1.55 pixels or 257 nm (continuous), 2.05 pixels or 340 nm (58 ms time-lapse), 2.92 pixels or 485 nm (200 ms time-lapse) and 3.62 pixels or 601 nm (1 s time-lapse). To make sure no freely diffusing molecules were tracked and to minimize false connections, a minimum track length of 5 frames was used for continuous and 3 frames for time-lapse movies. Other tracking settings were as described above.

Fluorescence survival time distributions of Squint and Cyclops were extracted from the single-molecule tracks, and GRID^52^ was used to determine the dissociation rate spectrum. Binding times were calculated as the inverse of the dissociation rate. The tails of the survival time distributions were cut off below a probability of 0.01 due to a low number of events.

The rate spectrum is a measure for how often dissociation events of a certain dissociation rate population occur within a specific time (Supplementary Figure 6). This can be converted into a “state” spectrum by dividing the fractions of the event spectrum with the respective dissociation rates. This results in the distribution of binding states at any given point in time. To estimate errors of the dissociation rate spectra, a 500× resampling was performed with randomly selected 80% of the data. Boundaries for dissociation rate clusters were then manually assigned to calculate the standard deviation of the results.

### Agent-based modelling

Agent-based models were implemented in Python3^82^. Morphogens were modelled as agents performing a random walk on a two-dimensional grid. The grid was generated in Fiji^83^ by scaling the binary masks of extracellular space such that each pixel in the image was 10 nm × 10 nm. Reflective boundary conditions were used. All intracellular pixels were set to 0 and extracellular to 1. The extracellular space was populated with 100 morphogens at random starting positions with a fraction of morphogens in bound state. The initial bound fraction (*BF*_*I*_) was estimated using an empirical function of the receptor density (σ), residence time (*τ*), and binding strength (*S*) to ensure that the equilibrium bound fractions were achieved quickly during the simulation.

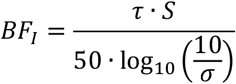

Morphogens were allowed to occupy only the extracellular positions. The receptors were placed at the boundary of the extracellular region with uniform spacing based on the required receptor density. The morphogen position and state (bound or unbound) was updated at each simulation step (10000 steps of 10 ms each). To simulate random walks of morphogens, their jump distances (*r*) were drawn from a range of jumps with probabilities given by the probability density function:

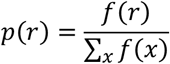

given that

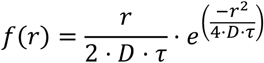

where, *D* is the diffusion coefficient (either *D*_*free*_ or *D*_*bound*_), *τ* is the timestep (10 ms), and *r, x* ∈ [0, 2].

The free and bound diffusion coefficients were set to 30 µm^2^/s (*D*_free_), and 0.5 µm^2^/s (*D*) as estimated from the experimental observations. A new position in the extracellular space was picked randomly from the available positions at distance *r* from the current position. The morphogen state was changed from unbound to bound if the distance to a receptor was ≤ 20 nm. The morphogen stayed bound for the time 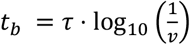, where *v* is a random number between 0 and 1, and *τ* is the residence time. The jump and angular histograms, and localization plots were generated in a similar manner as the experimental dataset to validate the model.

For the parameter screen, the two-dimensional grid was modified to narrow (*η* > 0) or widen (*η* < 0) the extracellular space. This was achieved by iteratively changing the pixel values at the extracellular boundary. For each iteration, the grid width was changed by 20 nm. The average width of the extracellular region was calculated to be approximately 1 µm, and *η* values ranging from -160 nm to 160 nm were tested. Three receptor densities σ (0.05, 0.5, 5.0 µm^-1^) and 5 residence times *τ* (0, 1. 6, 11. 16 s) were tested.

### Morphotrap experiments and image analysis

Confocal images of zebrafish embryos injected with different amounts of membrane-tethered GFP-binding nanobody were acquired as described previously^60^. Image analysis was performed manually in Fiji^83^. The GFP channel of the confocal images was converted into a .tif file. Five regions of interest (ROIs, 64×171 pixel) were manually selected from the image. For each ROI, 15 cavity and interface regions were selected, and the mean grey value was measured to quantify the GFP localization in each region. The ratio of the mean grey value of the cavity to that of the interface was used as ρ_c_/ρ_i_.

## Acknowledgements

We thank Gilbert Weidinger (Ulm University) for his constant support by providing access to his zebrafish facility, members of the Gebhardt and Michaelis labs for helpful discussions, Karlotta Bosch and Maximilian Haas for help cloning the HaloTag constructs, and Catrin Weiler for generating *oep* mRNA. The work was funded by the European Research Council (ERC) under the European Union’s Horizon 2020 Research and Innovation Program (No. 637987 ChromArch to J.C.M.G., No. 637840 QUANTPATTERN to P.M., No. 863952 ACE-OF-SPACE to P.M.) and the German Research Foundation (No. 422780363 SPP 2202 GE 2631/2–1 and No. 427512076 GE 2631/3–1 to J.C.M.G.). Support by the Collaborative Research Centre 1279, the Center for Translational Imaging MoMAN of Ulm University and the International Max Planck Research School “From Molecules to Organisms” is acknowledged.

## Contributors

P.M. and J.C.M.G. conceived the project; T.K., D.M., P.M. and J.C.M.G. designed the project; D.M. cloned morphogen fusion proteins and performed confocal microscopy; T.K. performed the measurements; J.C. contributed to the measurements; T.K. programmed TrackIt; J.G. and T.K. created the CNN segmentation tool; T.K. and J.C.M.G. analyzed data; A.N.L. performed simulations; T.K., A.N.L., P.M. and J.C.M.G wrote the manuscript with comments from all authors.

## Data availability

Data supporting the findings of this manuscript will be available from the corresponding authors after publication upon reasonable request. All single particle tracking data and simulated tracks will be freely available upon publication.

## Code availability

The TrackIt software is freely available. TrackIt was written in Matlab and is available at https://gitlab.com/GebhardtLab/TrackIt. The code for the agent-based model is available at https://github.com/mueller-lab/morphogenDiffusion-ABM.

