## Supplementary Figures and Tables for "Single-molecule tracking of Nodal and Lefty in live zebrafish embryos supports hindered diffusion model"

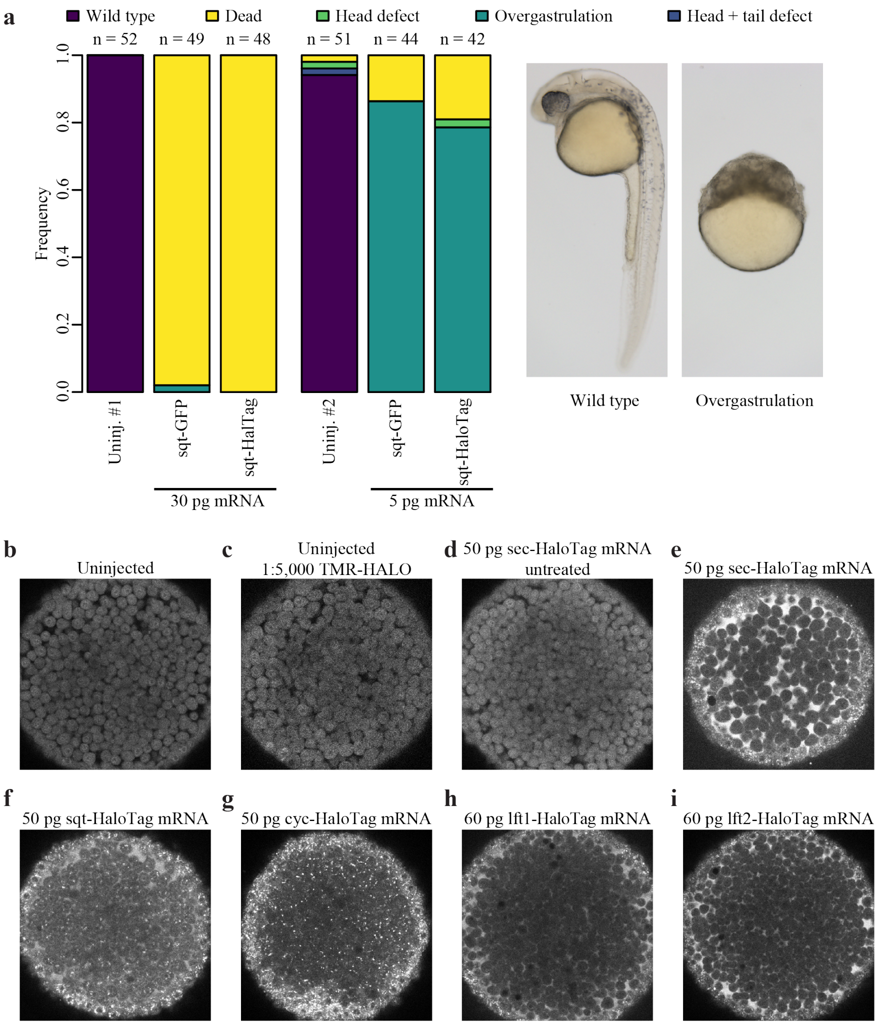

**Supplementary Figure 1. Activity and localization of HaloTag fusions. a)** The phenotypes upon injection of *squint-GFP* or *squint-HaloTag* mRNA are comparable for both high (30 pg mRNA, left) and low (5 pg mRNA, right) expression levels. **b-i)** Representative optical slices (animal views) of embryos that were uninjected (b), uninjected but treated with TMR-HALO ligand (c), injected with secreted-HaloTag-encoding mRNA but not treated with TMR-HALO ligand, and embryos expressing Secreted-HaloTag (e), Squint-HaloTag (f), Cyclops-HaloTag (g), Lefty1-HaloTag (h), or Lefty2-HaloTag (i) which were labeled with TMR-HALO ligand. Background fluorescence is similar in unlabeled Secreted-HaloTag-expressing embryos and TMR-HaloTag-treated uninjected embryos, but note that the fluorescence intensities between panels are not comparable due to different detector gain settings.

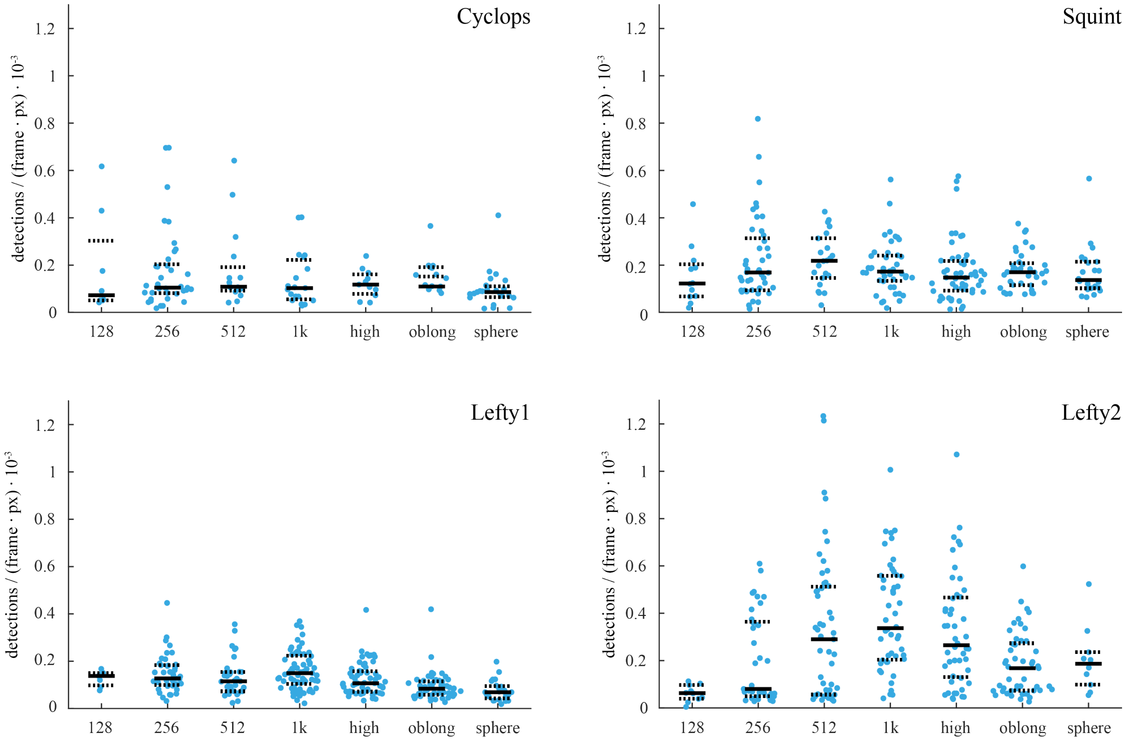

**Supplementary Figure 2. Average number of extracellular single molecule detections per frame in each movie.** Solid black lines indicate the median values, dashed black lines the 0.25 and 0.75 quantiles.

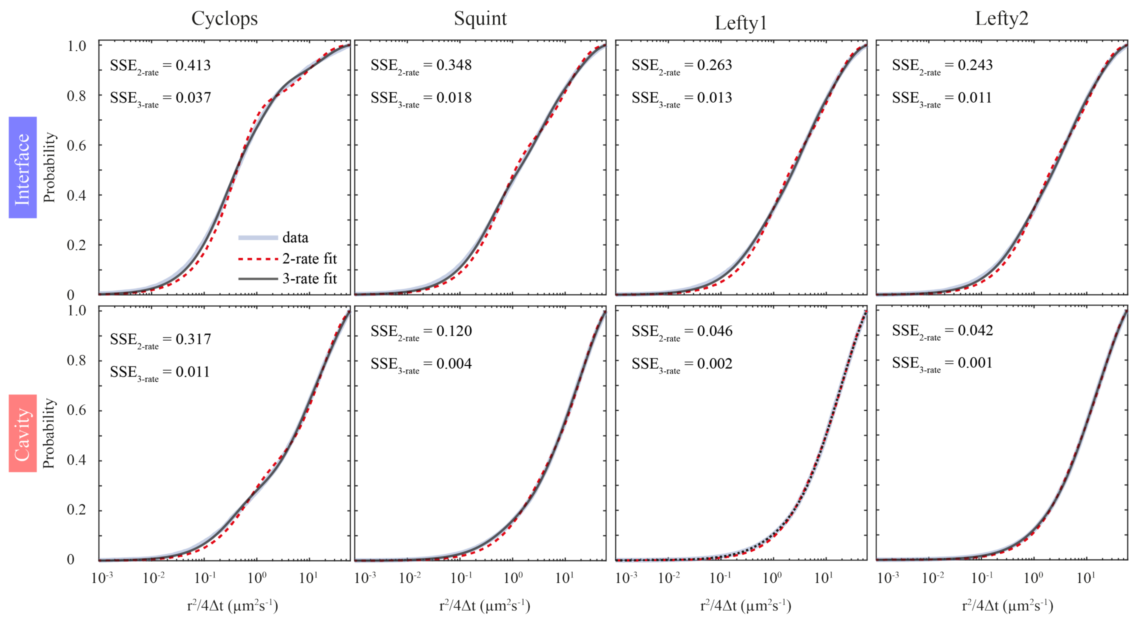

**Supplementary Figure 3**. **Fitting of jump distance distributions with a 2-rate and 3-rate Brownian diffusion model in interface (top) and cavity (bottom) regions for Cyclops, Squint, Lefty1 and Lefty2.** Based on the sum of squared errors (SSE), a three-component diffusion model best described the data.

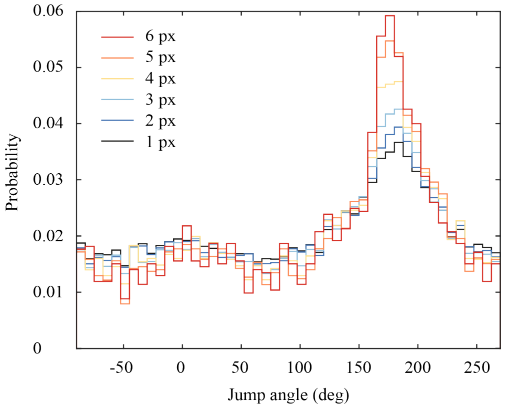

**Supplementary Figure 4. Jump angle distribution of Lefty2 in cavity regions as a function of the minimum distance of the jumps making up the angle.** The angle distributions exhibited a prominent contribution of reverse motion which increased with the minimum jump distance making up the angle, probably reflecting free diffusion in limited space in cavities.

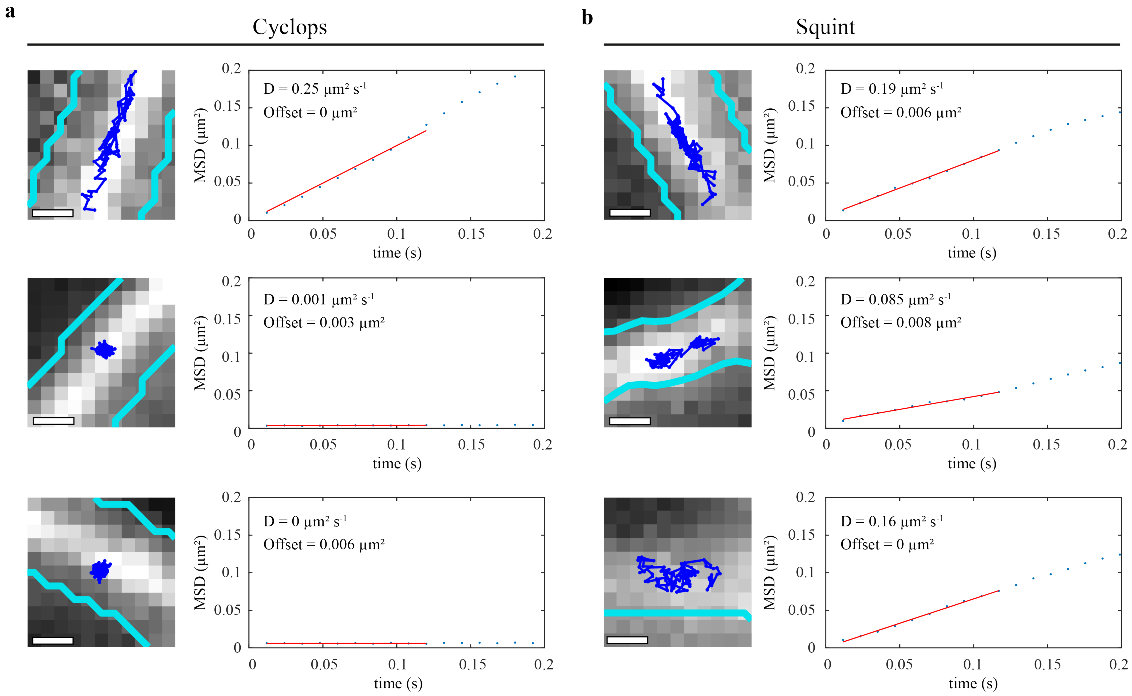

**Supplementary Figure 5. Mean-squared displacement (MSD) analysis of an example set of extracellular binding events of Cyclops and Squint (Supplementary Movie 6 and 7). a,b)** Left: memGFP signal averaged over 10 frames with overlaid track of a binding event (blue) with a minimum duration of 20 frames (234 ms) in the extracellular region (cyan). Right: MSD as a function of time (blue dots) with overlaid linear fit (red) for the determination of the diffusion coefficient of the tracks depicted on the left. Scale bars: 0.5 µm.

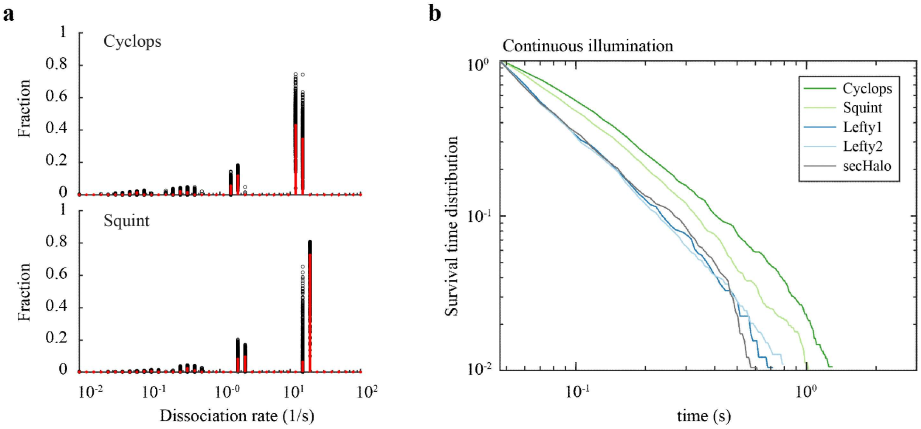

**Supplementary Figure 6. Residence times in the extracellular space. a)** Rate spectra of dissociation rates of Cyclops and Squint obtained by GRID using all data (red bars) and 500 resampling runs with randomly selected 80% of data (black data points) as an error estimation of the spectra. **b)** Survival-time distributions of bound molecules obtained from continuous movies.

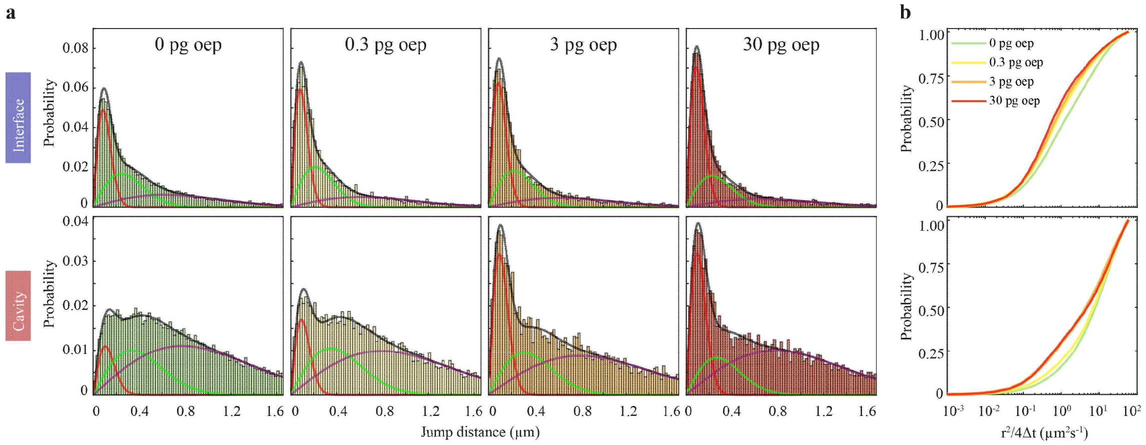

**Supplementary Figure 7. Mobility of Squint decreases with overexpression of oep. a)** Distribution of jump distances within single-molecule tracks in interfaces and cavities of Squint with the indicated amount of co-injected *oep*. Lines represent a three-component diffusion model (black) and the individual components (red, green, purple). **b)** Cumulative distributions of jump distances.

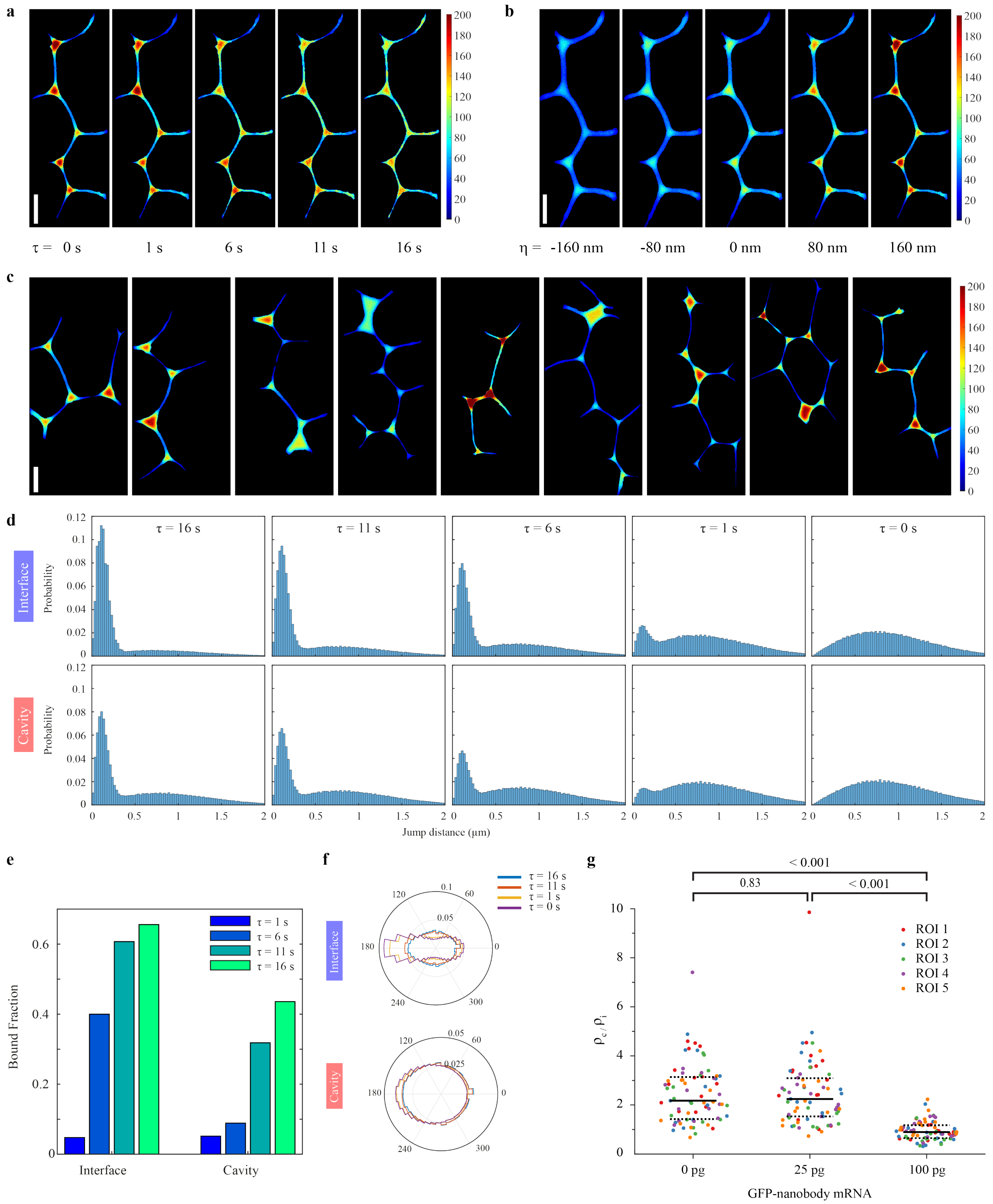

**Supplementary Figure 8. An agent-based model reveals key parameters affecting morphogen behavior in the extracellular space. a) a)** Localization density plots of simulated morphogens with increasing residence times ($\tau$) at $\sigma$ =0.5 µm$^{-1}$ and $\eta$ = 160 nm. **b)** Localization density plots of simulated morphogens with increasing narrowness ($\eta$) at $\tau$ = 0 s, and $\sigma$ = 0.5 µm$^{-1}$. **c)** Localization density plots of morphogens simulated on different grids; $\eta$ = 160 nm, $\tau$ = 0 s, and $\sigma$ = 0.5 µm$^{-1}$ **d)** Distribution of jump distances of simulated morphogens in interfaces and cavities for different residence times. **e)** Fraction of bound molecules in interfaces and cavities for different residence times. **f)** Distributions of angles between consecutive track segments of simulated morphogens in interfaces and cavities for different residence times. **g)** Ratio of mean GFP intensities in cavity and interface ($\rho_c/\rho_i$) for zebrafish embryos injected with different amounts of mRNA encoding membrane-tethered GFP-binding nanobody. Solid black lines indicate the median values, dashed black lines the 0.25 and 0.75 quantiles..

The scattered dots show individual data points and are color-coded based on the regions of interest (ROIs) used for measurements. For each condition, fifteen cavity and interface regions each were measured from five ROIs. P-values were calculated using the Kruskal-Wallis-Test. Scale bars in a, b and c are 10 µm.

### Supplementary Movie Legends

**Supplementary Movie 1. Visualization of the data demonstrated in Figure 1.** Left: single molecule imaging of Lefty2-Halo molecules imaged with 11.7 ms per frame. Center: memGFP signal averaged over 10 frames with overlaid interface (blue) and cavity (red) regions and tracked single molecules Right: TALM image showing the amount of detections in each pixel over the movie time. Scale bar: 10 µm.

**Supplementary Movie 2-5. Visualization of the data demonstrated in Figure 2.** Left: single molecule imaging of Halo-Cyclops, Halo-Squint, Lefty1-Halo and Lefty2-Halo at 11.7 ms per frame. Right: TALM image showing the amount of detections in each pixel over the movie time with overlaid interface (blue) and cavity (red) regions and tracked single molecules. Scale bar: 5 µm.

**Supplementary Movie 6-7. Visualization of the data demonstrated in Supplementary Figure 5.** Top: single molecule imaging of Halo-Cyclops and Halo-Squint at 11.7 ms per frame. Bottom: memGFP signal averaged over 10 frames with overlaid extracellular region (cyan) and track of a binding event. Scale bar: 0.5 µm.

### Supplementary Tables

| **Interface** | Cyclops | Squint | Lefty1 | Lefty2 | secHalo |
|---|---|---|---|---|---|
| $D_1$ ($\mu m^2 s^{-1}$) | 0.2 ± 0.02 | 0.32 ± 0.01 | 0.36 ± 0.04 | 0.44 ± 0.03 | 0.47 ± 0.04 |
| $D_2$ ($\mu m^2 s^{-1}$) | 1.16 ± 0.11 | 2.61 ± 0.2 | 2.65 ± 0.25 | 3.04 ± 0.22 | 3.84 ± 0.29 |
| $D_3$ ($\mu m^2 s^{-1}$) | 16.66 ± 1.13 | 16.59 ± 0.76 | 16.05 ± 0.83 | 17.02 ± 0.68 | 23.82 ± 1.19 |
| $A_1$, immobile | 0.44 ± 0.03 | 0.35 ± 0.01 | 0.21 ± 0.02 | 0.24 ± 0.01 | 0.17 ± 0.01 |
| $A_2$, intermediate | 0.39 ± 0.03 | 0.34 ± 0.01 | 0.39 ± 0.01 | 0.39 ± 0.01 | 0.38 ± 0.01 |
| $A_3$, fast | 0.17 ± 0.01 | 0.31 ± 0.01 | 0.4 ± 0.02 | 0.37 ± 0.02 | 0.45 ± 0.02 |

**Supplementary Table 1.** Diffusion parameters in interface regions obtained from fitting the cumulative distribution of jump distances of Cyclops, Squint, Lefty1 and Lefty2 shown in **Error! Reference source not found.**.

| **Cavity** | Cyclops | Squint | Lefty1 | Lefty2 | secHalo |
|---|---|---|---|---|---|
| $D_1$ ($\mu m^2 s^{-1}$) | 0.31 ± 0.02 | 0.45 ± 0.04 | 0.64 ± 0.11 | 0.73 ± 0.09 | 1.16 ± 0.15 |
| $D_2$ ($\mu m^2 s^{-1}$) | 5.19 ± 0.73 | 4.84 ± 0.44 | 6.8 ± 0.72 | 5.54 ± 0.37 | 8.04 ± 0.8 |
| $D_3$ ($\mu m^2 s^{-1}$) | 28.03 ± 3.22 | 25.82 ± 1.11 | 30.05 ± 2.09 | 26.83 ± 1.01 | 38.96 ± 2.81 |
| $A_1$, immobile | 0.22 ± 0.01 | 0.09 ± 0.01 | 0.05 ± 0.01 | 0.06 ± 0.01 | 0.06 ± 0.01 |
| $A_2$, intermediate | 0.3 ± 0.04 | 0.28 ± 0.02 | 0.32 ± 0.03 | 0.33 ± 0.02 | 0.32 ± 0.03 |
| $A_3$, fast | 0.48 ± 0.04 | 0.63 ± 0.02 | 0.63 ± 0.04 | 0.61 ± 0.02 | 0.61 ± 0.03 |

**Supplementary Table 2.** Diffusion parameters in cavity regions obtained from fitting the cumulative distribution of jump distances of Cyclops, Squint, Lefty1 and Lefty2 shown in **Error! Reference source not found.**.

| **Interface** | *oep* 0.3pg | *oep* 3pg | *oep* 30pg |
|---|---|---|---|
| $D_1$ ($\mu m^2 s^{-1}$) | 0.28 ± 0.02 | 0.31 ± 0.02 | 0.32 ± 0.02 |
| $D_2$ ($\mu m^2 s^{-1}$) | 1.9 ± 0.18 | 2.07 ± 0.29 | 2.18 ± 0.27 |
| $D_3$ ($\mu m^2 s^{-1}$) | 15.63 ± 0.9 | 16.44 ± 1.34 | 17.82 ± 1.46 |
| $A_1$, immobile | 0.4 ± 0.02 | 0.44 ± 0.03 | 0.5 ± 0.02 |
| $A_2$, intermediate | 0.35 ± 0.016 | 0.33 ± 0.018 | 0.29 ± 0.016 |
| $A_3$, fast | 0.25 ± 0.013 | 0.23 ± 0.017 | 0.2 ± 0.014 |

**Supplementary Table 3.** Diffusion parameters in interface regions obtained from fitting the cumulative distribution of jump distances of Squint upon overexpression of *oep* with 0.3 pg, 3 pg and 30 pg mRNA shown in **Error! Reference source not found.**.

| **Cavity** | *oep* 0.3pg | *oep* 3pg | *oep* 30pg |
|---|---|---|---|
| $D_1$ ($\mu m^2 s^{-1}$) | 0.37 ± 0.03 | 0.38 ± 0.03 | 0.38 ± 0.04 |
| $D_2$ ($\mu m^2 s^{-1}$) | 5.18 ± 0.57 | 4.22 ± 0.9 | 3.23 ± 1.08 |
| $D_3$ ($\mu m^2 s^{-1}$) | 27.1 ± 1.93 | 28.09 ± 3.58 | 26.33 ± 2.62 |
| $A_1$, immobile | 0.13 ± 0.01 | 0.25 ± 0.02 | 0.25 ± 0.03 |
| $A_2$, intermediate | 0.3 ± 0.03 | 0.25 ± 0.03 | 0.19 ± 0.02 |
| $A_3$, fast | 0.57 ± 0.03 | 0.5 ± 0.04 | 0.56 ± 0.03 |

**Supplementary Table 4.** Diffusion parameters in cavity regions obtained from fitting the cumulative distribution of jump distances of Squint with overexpression of *oep* with 0.3 pg, 3 pg and 30 pg mRNA shown in **Error! Reference source not found.**

| | Measurement days | Movies | Tracks | | | Jumps | | | |
|---|---|---|---|---|---|---|---|---|---|
| | | | Interface | Cavity | Combined | Interface | Cavity | Combined | Jumps/movie |
| Cyclops | 7* | 128 | 2837 | 4063 | 6900 | 8569 | 8685 | 17254 | 135.5 |
| Squint | 10* | 231 | 6903 | 15842 | 22745 | 31615 | 17983 | 49598 | 214.7 |
| Lefty1 | 4 | 289 | 5012 | 18474 | 23486 | 32295 | 11050 | 43345 | 150.1 |
| Lefty2 | 4 | 253 | 9365 | 28642 | 38007 | 62118 | 21539 | 83657 | 330.6 |
| Squint + 0.3 pg oep | 2 | 200 | 2987 | 6425 | 9412 | 12495 | 7070 | 19565 | 103.6 |
| Squint + 3 pg oep | 3 | 288 | 2875 | 3589 | 6464 | 6917 | 7721 | 14638 | 50.8 |
| Squint + 30 pg oep | 3 | 255 | 2729 | 3067 | 5796 | 5533 | 8224 | 13757 | 49.4 |
| secHalo | 3 | 281 | 9969 | 28655 | 38624 | 20585 | 52351 | 72936 | 259.6 |

**Supplementary Table 5.** Statistics for continuous movie analysis. *Total number of measurement days for continuous and time-lapse movies.

| | Movies | Tracks | Tracks per movie |
|---|---|---|---|
| Cyclops 12 ms | 128 | 1418 | 11.078125 |
| Squint 12 ms | 252 | 1782 | 7.071428571 |
| Cyclops 58 ms | 78 | 883 | 11.32051282 |
| Squint 58ms | 131 | 886 | 6.763358779 |
| Cyclops 200 ms | 91 | 986 | 10.83516484 |
| Squint 200 ms | 135 | 770 | 5.703703704 |
| Cyclops 1000 ms | 93 | 550 | 5.913978495 |
| Squint 1000 ms | 119 | 508 | 4.268907563 |

**Supplementary Table 6.** Statistics of single-molecule tracks.
